# The *C.elegans* AWA Olfactory Neuron Fires Calcium-Mediated All-or-None Action Potentials

**DOI:** 10.1101/359935

**Authors:** Qiang Liu, Philip B. Kidd, May Dobosiewicz, Cornelia I. Bargmann

## Abstract

We find, unexpectedly, that *C. elegans* neurons can encode information through regenerative all-or-none action potentials. In a survey of current-voltage relationships in *C. elegans* neurons, we discovered that AWA olfactory neurons generate membrane potential spikes with defining characteristics of action potentials. Ion substitution experiments, pharmacology, and mutant analysis identified a voltage-gated CaV1 calcium channel and a Shaker-type potassium channel that underlie action potential dynamics in AWA. Simultaneous patch-clamp recording and calcium imaging in AWA revealed spike-associated calcium signals that were also observed after odor stimulation of intact animals, suggesting that natural odor stimuli induce AWA action potentials. The stimulus regimes that elicited action potentials match AWA’s proposed specialized function in climbing odor gradients. Our results provide evidence that *C. elegans* can use digital as well as analog coding schemes, expand the computational repertoire of its nervous system, and inform future modeling of its neural coding and network dynamics.

## Introduction

The nervous system uses both digital and analog signals to encode perception, computation, and action (Rieke et al., 1997; Dayan and Abbott, 2005). Spiking neurons, which are prominent in most animal nervous systems, compress continuous inputs into digital, regenerative action potentials that transmit information efficiently over long distances. Graded neurons can encode more than one bit of information per time interval, but are more sensitive to noise than spiking neurons, and therefore are typically found in settings such as the vertebrate retina in which signals are propagated only over short distances (Koch et al., 2006; Sarpeshkar, 1998). Graded signaling allows parallel computations to occur in different parts of a neuron, and even spiking neurons use this multiplexing property in their dendritic arbors.

Among animals, nematodes have been thought to be exceptional in their lack of neuronal action potentials. Early recordings in the parasitic nematode *Ascaris suum* demonstrated only graded electrical properties and graded synaptic transmission in motor neurons (Davis and Stretton, 1989b), together with high membrane resistance that allows these neurons to propagate signals over centimeters without action potentials (Davis and Stretton, 1989a). A variety of graded electrical properties have subsequently been described in the free-living nematode *Caenorhabditis elegans* (Geffeney et al., 2011; Goodman et al., 1998; Lindsay et al., 2011; Liu et al., 2017; Liu et al., 2014; Liu et al., 2009; Mellem et al., 2008; O’Hagan et al., 2005; Ramot et al., 2008). Some neurons are smoothly depolarized or hyperpolarized from a tonic resting potential (Liu et al., 2014; Mellem et al., 2008), while others are bistable, with nonlinear transitions between the resting potential and a depolarized potential around −20 mV (Goodman et al., 1998). Striking nonlinearity is observed in the regenerative plateau potentials of RMD motor neurons, which resemble action potentials in their all-or-none properties, but do not spontaneously repolarize (Mellem et al., 2008).

The apparent absence of canonical action potentials in nematode neurons aligns with the absence of genes encoding voltage-gated sodium channels in the *C. elegans* genome (Bargmann, 1998). However, voltage-gated calcium channels can generate alternative modes of spiking: for example, dendritic calcium spikes play important integrative roles in vertebrate pyramidal neurons (Golding et al., 1999; Larkum et al., 1999). Calcium-dependent action potentials at 3-10 Hz have been recorded from *C. elegans* pharyngeal muscles (Raizen and Avery, 1994; Shtonda and Avery, 2005) and body-wall muscles (Gao and Zhen, 2011; Liu et al., 2011). Activity patterns suggestive of neuronal calcium spikes have been observed in extracellular recordings in the *Ascaris* ventral nerve cord (Davis and Stretton, 1992), and in all-or-none small-amplitude spikes in a *C. elegans* interneuron (Faumont et al., 2012), suggesting that further investigation of such events could be fruitful.

There are only 302 neurons in the adult *C. elegans* hermaphrodite, but their functions, anatomy, molecular properties, and synaptic partners allow them to be divided into at least 120 classes (White et al., 1986). The preferential use of analog over digital signaling in *C. elegans* neurons increases the theoretical coding capacity of its tiny neuronal repertoire. At the same time, some neurons among these diverse types might prefer digital regimes to engage the coding properties ascribed to spike rates or spike timing (Gerstner and Kistler, 2002). We show here that AWA olfactory neurons, which are specialized for detecting gradients of attractive odors, reliably and reproducibly generate all-or-none action potentials under physiologically realistic conditions, and explore the mechanisms and implications of this property.

## Results

### AWA neurons fire all-or-none action potentials

To explore the biophysical diversity of the *C. elegans* nervous system, we conducted an electrophysiological survey of identified *C. elegans* head neurons. At least four distinct types of current-voltage (I-V) relationships were observed (Fig 1). A motor-neuron/interneuron, RIM, exhibited near-zero inward currents under hyperpolarizing voltage-clamp steps and rapid inactivating outward currents under depolarizing steps (Fig 1B). The lack of large sustained currents allowed RIM to be depolarized or hyperpolarized smoothly under current-clamp (Fig 1A). We defined neurons with this type of I-V relationship as “transient outward rectifiers”. The interneuron AIY also lacked large inward currents under hyperpolarizing steps, but had non-inactivating outward currents under depolarizing steps (Fig 1B), as previously reported (Faumont et al., 2006). These outward currents made AIY’s membrane potential more sensitive to hyperpolarizing than depolarizing inputs with a transition point around −30 mV (Fig 1C). Neurons with this type of I-V relationship were defined as “outward rectifiers”. The third type of neuron, “bistable”, was inwardly and outwardly rectifying, and included AFD thermosensory neurons (Fig 1B). In agreement with a previous study (Ramot et al., 2008), large sustained currents to hyperpolarizing and depolarizing steps in AFD essentially restricted its membrane potential to two voltages around −80 mV and −20 mV over a range of current injections (Fig 1A). Finally, like AFD, the AWA olfactory neuron had large sustained outwardly- and inwardly-rectifying currents in response to depolarizing and hyperpolarizing steps (Fig 1B C). In addition, and to our surprise, AWA membrane potential exhibited oscillations reminiscent of classic Hodgkin– Huxley action potentials under stimulations of certain amplitudes and durations in current clamp (Fig 1A). We define AWA as a “bistable/spiking” neuron.

**Figure 1.**
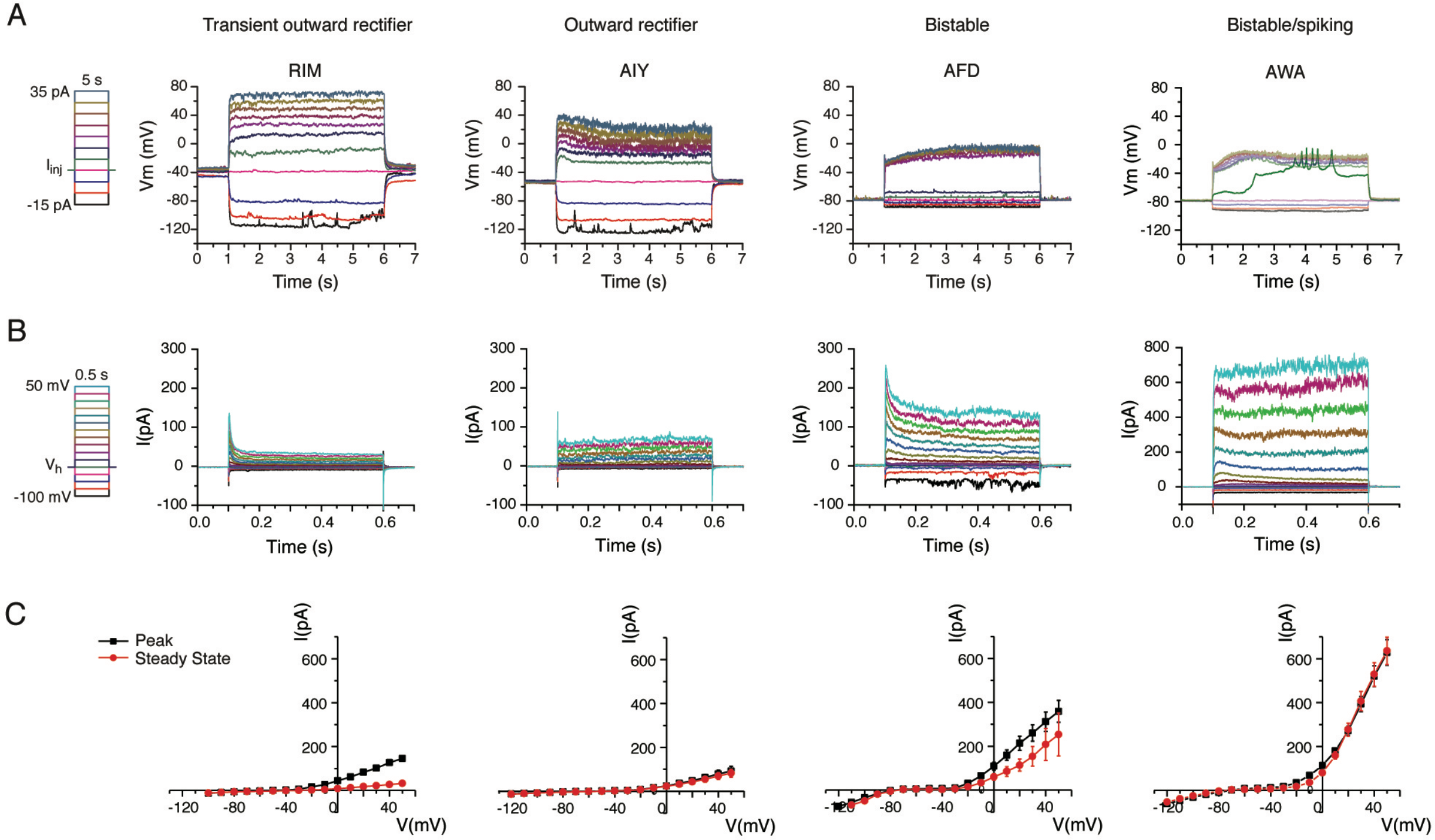
Electrophysiological diversity of C. elegans neurons. **A.** Examples of membrane potential dynamics induced by a series of current injection steps in current-clamped RIM, AIY, AFD and AWA neurons. Left, current injection protocol, starting from −15 pA and increasing to 35 pA by 5 pA increments. For AWA, the green membrane potential trace with spiking activities was triggered by the 5 pA current injection step. Non-spiking traces were dimmed for clarity. **B.** Examples of whole-cell current traces induced by a series of voltage steps in voltage-clamped RIM, AIY, AFD and AWA (Note difference in Y axis in AWA neurons). Left, voltage step protocol, starting from −100 mV and increasing to 50 mV in 10 mV increments. The holding potential was set at −60 mV before and after applying each voltage step. **C.** I-V relationships for the four neurons above obtained from averaged voltage-clamp recordings. Black squares: peak currents measured by the absolute maximum amplitude of currents within first 100 ms of each voltage step onset. Red circles: steady-state currents measured by the averaged currents of the last 50 ms of each voltage step. Number of cells tested for each neuron: RIM: n = 3; AIY: n = 7; AFD: n = 3; AWA: n = 16. Error bars show standard error of the mean (SEM).

AWA spikes could occur either as a single spike or in a transient burst of spikes at up to 10.5 Hz (mean frequency: 4.2 ± 0.2 Hz, n = 89; error denotes SEM in all cases) (Fig 2A, I, Fig S1A, C-E). All AWA spikes had a stereotypical shape with a fast upstroke followed by a fast downstroke and afterhyperpolarization (AHP). Spikes were all-or-none with an amplitude that was well-fit by a normal distribution (Fig 2B-D). Single spikes had a threshold of −34.0 ± 0.6 mV, an amplitude of 25.8 ± 0.8 mV, a half-maximum width of 18.0 ± 0.8 ms, and an AHP of 8.4 ± 0.9 mV (Fig 2C). The presence of adjacent spikes during bursts altered the membrane potential before the spike and the magnitude of the AHP (Fig 2B, Table S3). The stereotypical shape of single or bursting spikes was invariant with respect to the amplitude, duration, and waveform of the input stimulus. We conclude that AWA fires all-or-none action potentials.

**Figure 2.**
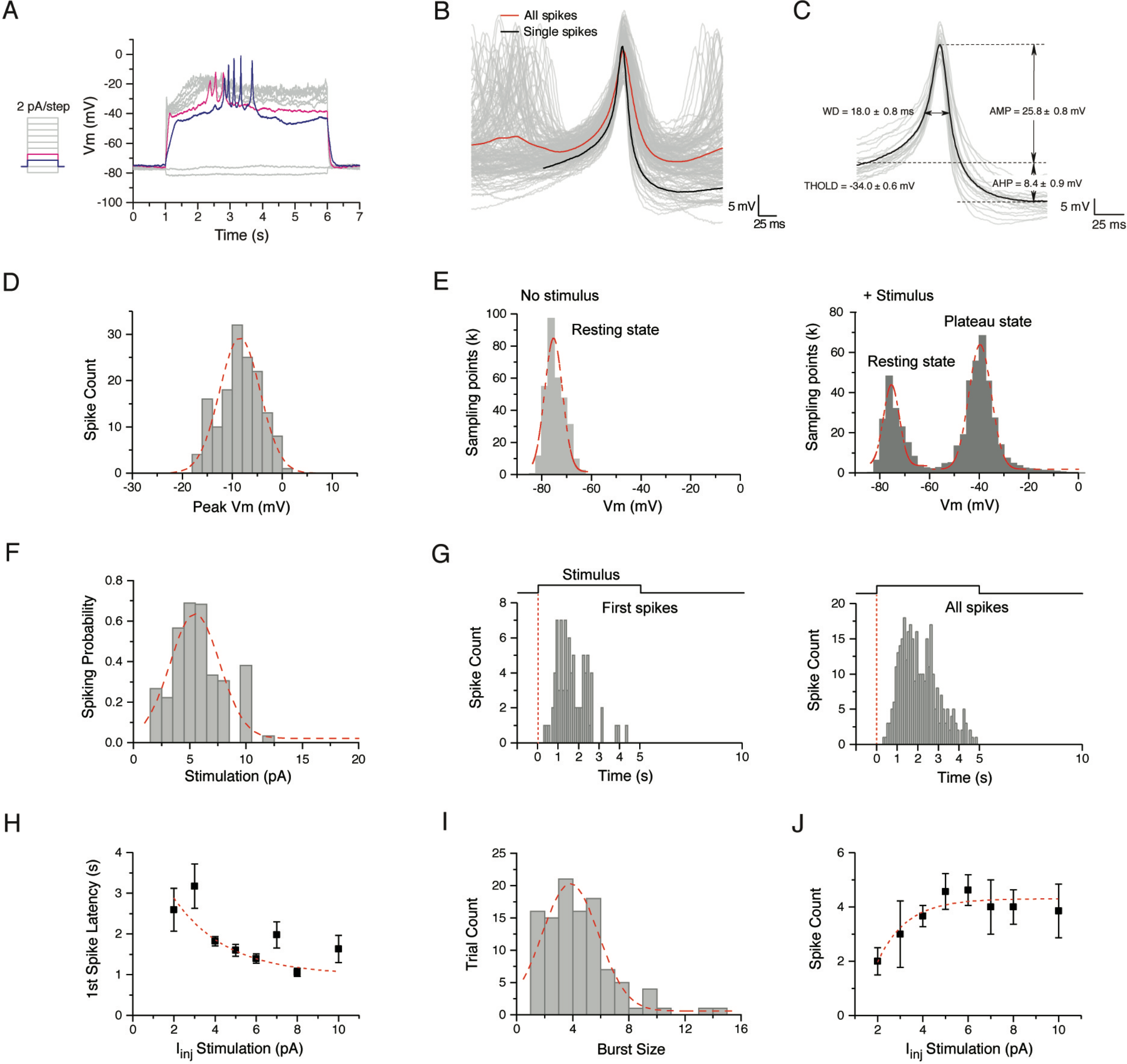
AWA neurons fire all-or-none action potential. **A.** Representative current-clamp recording traces in AWA under a series of 2 pA-increment current injection steps. The current injection protocol used is on the left (same below and in all figures). **B.** Average action potential trace from 19 single spikes (black) and average over all 149 spikes including those in bursts (red). **C.** Statistics of the average action potential trace (black) from 19 single spikes. THOLD: threshold potential; AMP: amplitude relative to threshold; WD: half-maximum width; AHP: afterhyperpolarization. Numbers shown as mean ± SEM. Statistics across all 149 spikes are provided in Table S3. In **B** and **C**, Individual traces aligned by the maximum slope of the upstroke are in grey. **D.** Amplitude histogram of peak membrane potential of the 149 action potential spikes in **B**. Bin size 2 mV. Mean = −8.1 mV, SEM = 0.6 mV, R-Square = 0.84. **E.** Left: AWA resting membrane potential in 27 traces that yielded action potentials (shown in Fig S1D). Mean = −75.3 mV, SEM = 4.7 mV, R-Square = 0.90. Right: after current injection, AWA membrane potential resolves into two peaks: (1) Resting state: Mean = −75.4 mV, SEM = 4.2 mV, R-Square = 0.91; (2) Plateau state: Mean = −39.6 mV, SEM = 1.4 mV, R-Square = 0.99. **F.** Spiking probability, calculated from the number of trials when spikes were successfully induced by a specific current injection step divided by the number of total trials when such steps were used. Mean = 5.5 pA, SEM = 0.3 pA, R-Square = 0.84, n = 119 spikes. Spiking was only induced by current steps between +2 and +12 pA; tested range was −10 to +35 pA in 1,2, or 5 pA increments (n=17-56 cells). **G.** Spike time distribution of first spike (left) and all spikes (right) under 5 s of current injection steps at all amplitudes. Bin size is 100 ms. Note delay after the stimulus onset before spiking. **H.** First-spike latency decreases with injected current amplitude. Red dotted line is fitted with an exponential decay function (R-Square = 0.72). **I.** Burst size (spike count in a burst) of all spikes recorded in WT AWA under 5 s of current injection steps at all amplitudes. Red dotted line is fitted with a Gaussian distribution function (Mean = 3.8, SEM = 0.2, R-Square = 0.89). **J.** Burst size increases moderately with injected current amplitude. Red dotted line is fitted with exponential decay function (R-Square = 0.89). Red dotted lines in **D-F** and **J** are fitted with a Gaussian distribution function.

In addition to these features common to all action potentials, action potentials in AWA had several characteristic features. First, they were associated with membrane potential bistability. Due to large depolarizing and hyperpolarizing currents after voltage steps, the AWA membrane potential had two stable states: a resting state at around −75 mV and a more depolarized plateau state at around −40 mV, near the firing threshold (Fig 2E). A small positive current caused a membrane potential jump from the resting state to the plateau state, but larger current injections resulted in little further depolarization unless spikes occurred. Second, action potentials were elicited only by stimuli of amplitudes between 2 pA and 12 pA, with the optimal 5 or 6 pA stimulus resulting in a ~70% spiking rate; no spiking was observed at higher current injections despite depolarization to the plateau state (Fig 2A, F). Third, spikes were induced only after a sustained current injection, with a peak time to first spike of one second (Fig 2G, Fig S1B-D). The latency to first spike decreased with increasing stimulus strength within the spike-generating range (Fig 2H). Fourth, bursts of spikes typically terminated after 1 - 8 spikes, with a total burst duration of less than 2 seconds, so that most spikes occurred between 1 - 4 seconds after stimulation (Fig 2G, I). Even a 30 second stimulus of appropriate amplitude only induced spikes within the first few seconds (Fig S1A). The exact timing and number of spikes induced by a given stimulus varied; for example, a square stimulus of the same amplitude and duration could induce a single spike or a burst after a variable delay (Fig S1E). There was a weak increase in spike count with stimulus strength (Fig 2J). These collective properties were reproducible across animals, and defined a narrow but reliable spiking regime (see Discussion).

### The CaV1-type calcium channel EGL-19 generates the action potential upstroke

We used genetic mutations, ion substitution, and pharmacological experiments to define the requirements for AWA spiking. AWA spiking was normal in *unc-13* mutants, which lack most fast synaptic transmission, suggesting that spiking results from intrinsic AWA biophysical properties (Fig 3A). Ionic substitution of extracellular Na^+^ with N-Methyl-D-glucamine (NMDG) during recording altered resting membrane potential but spared AWA action potentials (Fig 3B). By contrast, reducing extracellular Ca^2+^ to 0-1 mM eliminated AWA spikes (Fig 3C, Table S2). Conversely, increasing extracellular Ca^2+^ to 5 mM or 8 mM generated action potential spikes with larger amplitudes (Fig 3D, Fig S2A, Table S3). These results strongly suggest that action potentials in AWA are calcium-mediated.

**Figure 3.**
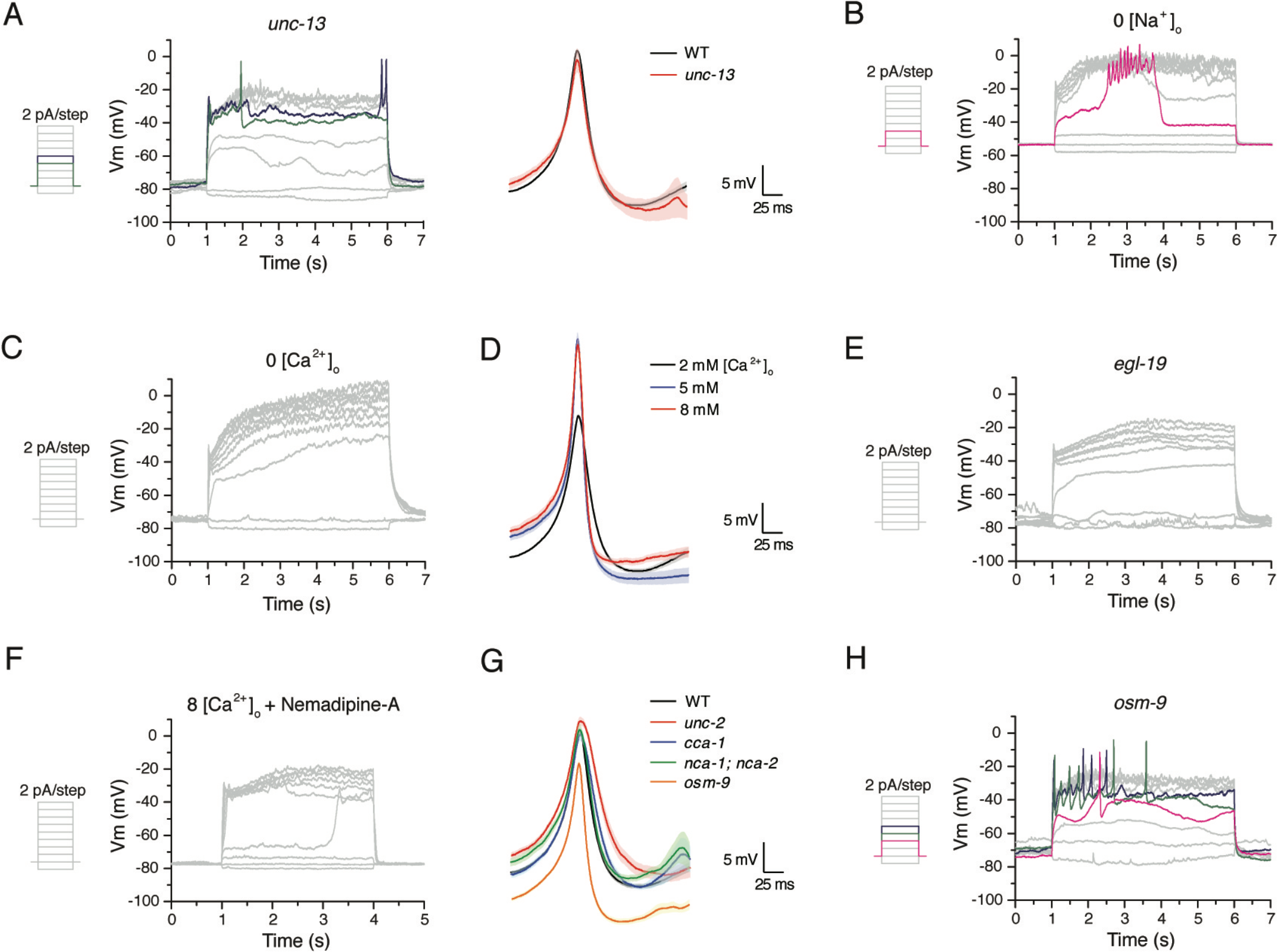
The CaV1-type calcium channel EGL-19 generates the action potential upstroke. **A.** Representative current-clamp traces recorded from *unc-13* AWA in 2 mM [Ca^2+^]_o_ extracellular solution (default [Ca^2+^]_o_ if not labeled otherwise). Insert: comparison of averaged action potential spikes from *unc-13* (n = 12) and WT (same as the red trace in Fig 2B). The shades are SEM (same below and in all figures). **B.** Representative current-clamp traces recorded from WT AWA in extracellular solutions with Na^+^ substituted by NMDG. **C.** Representative current-clamp traces recorded from WT AWA in extracellular solutions with 0 mM [Ca^2+^]_o_. **D.** Comparison of averaged action potential spikes from WT AWA in 2 (n = 19), 5 (n = 20) or 8 mM (n = 13) [Ca^2+^]_o_. The 2 mM [Ca^2+^]_o_ trace is the same as shown in Fig 2B. **E.** Representative traces recorded from *egl-19(n582)* AWA. **F.** Representative traces recorded from WT AWA with 10 uM Nemadipine-A and 8 mM Ca^2+^ in the extracellular solution. No action potentials were detected in **E** and **F**. **G.** Comparison of averaged action potential spikes from WT (same as the red trace in Fig 2B), *unc-2* (n = 13), *cca-1* (n = 18), *nca-1; nca-2* (n = 24) and *osm-9* (n = 55) AWAs. **H.** Representative current-clamp traces recorded from *osm-9* AWA. AWA spikes in calcium substitution experiments and in *osm-9* mutants are significantly different from WT (Table S3).

To identify channels responsible for the action potential in AWA, we recorded from mutant strains lacking each of the five predicted alpha subunits in the voltage-gated calcium channel family. AWA spikes could not be detected in reduction-of-function mutants in *egl-19*, which encodes a CaV1 L-type voltage-gated calcium channel (Fig 3E). In agreement with this result, acute addition of the selective L-type channel blocker Nemadipine-A blocked spiking in 6 out of 7 AWA neurons (Fig 3F). AWA spikes were normal in mutants for *unc-2* (CaV2 P/Q-type high voltage-gated calcium channel), *cca-1* (CaV3 T-type low voltage-gated calcium channel) (Frokjaer-Jensen et al., 2006) and *nca-1; nca-2* (NALCN-related voltage-independent, nonselective cation channels) (Jospin et al., 2007) (Fig 3G, Fig S2B, Table S3). The TRPV sensory channel OSM-9, which is expressed in AWA, has also been suggested to be voltage-activated (Colbert et al., 1997; Glauser et al., 2011), but action potentials were spared in *osm-9* mutants (Fig 3H) albeit with a shifted threshold suggesting a role for *osm-9* in AWA excitability (Fig 3G, Table S3). Together, these results point to a uniquely important role for EGL-19, a CaV1 type voltage-gated calcium channel, in AWA action potential generation.

### Shaker-type potassium channels drive the action potential downstroke

Termination of the action potential in the Hodgkin-Huxley model is a result of voltage-dependent potassium channels activated by depolarization. Eliminating potassium currents by substituting intracellular K^+^ with Cs^+^ and adding the potassium channel blockers tetraethylammonium (TEA) and 4-Aminopyridine (4-AP) to the extracellular solution resulted in a more linear AWA membrane potential without spiking, suggesting that potassium channels dominate AWA voltage dynamics inclusive of action potentials (Fig 4A).

**Figure 4.**
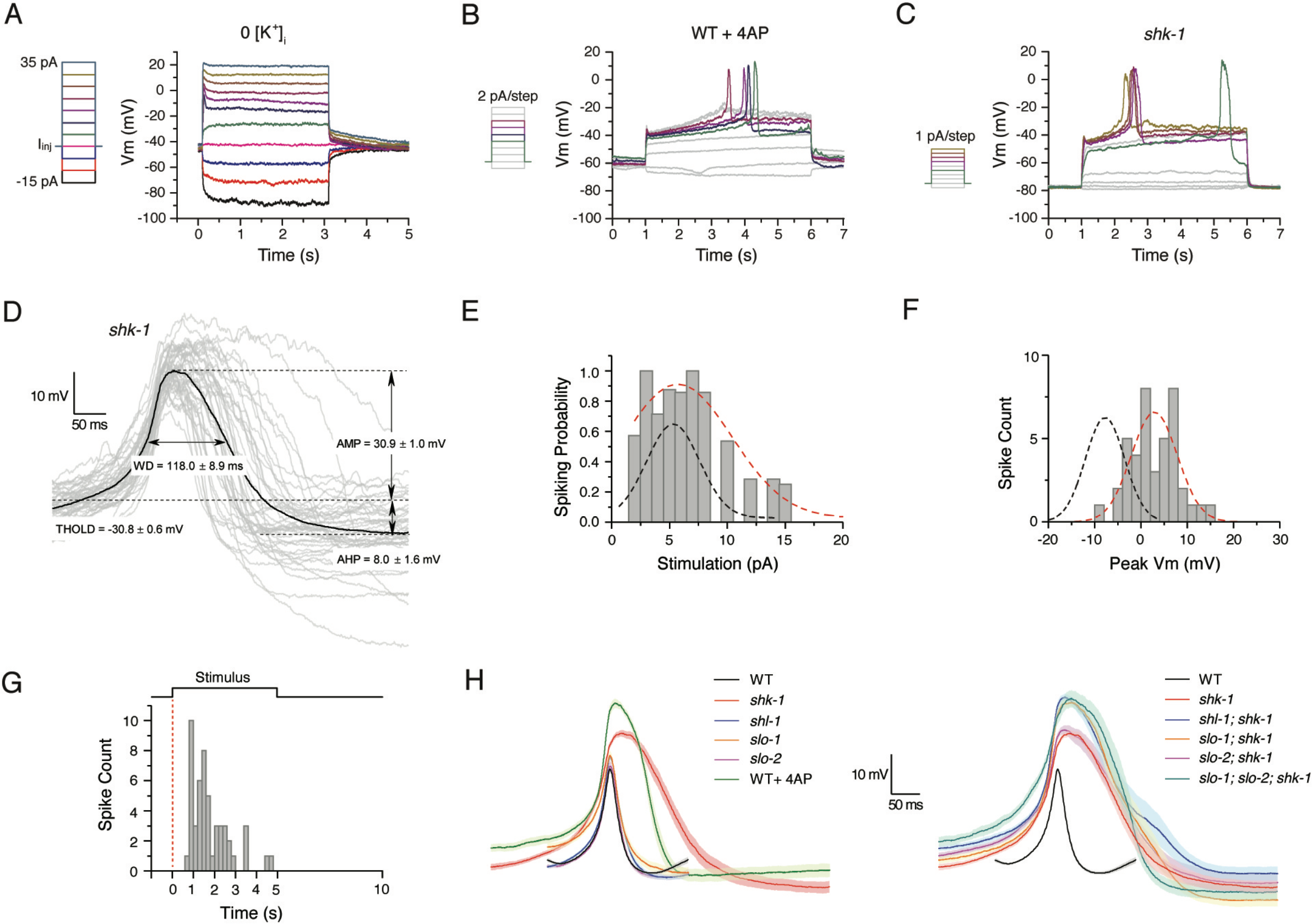
Shaker-type potassium channels drive the action potential downstroke. **A.** Representative current-clamp traces in AWA recorded with 0 mM K^+^ intracellular solution and 150 mM TEA and 3 mM 4-AP extracellular solution. Both bistability and spiking activity were absent. **B**. Representative current-clamp traces in AWA recorded in physiological solutions with 5 mM 4-AP. **C.** Representative current-clamp traces recorded in AWA in *shk-1*. **D.** Statistics of the averaged action potential trace from *shk-1* AWA (n = 41). The individual action potential traces used for averaging are shown in grey. Compare WT in panel **H** and Fig 2C. **E.** Spiking probability of *shk-1* AWA under various current injection amplitudes (Mean = 5.7 pA, SEM = 0.5 pA, R-Square = 0.91, n = 12). **F.** Amplitude histogram of all *shk-1* AWA action potential spikes used in **D** (Mean = 2.7 mV, SEM = 1.1 mV, R-Square = 0.60). Bin size is 2 mV. Red dotted lines in **E** and **F** are fitted with a Gaussian distribution function. Black dotted lines in **E** and **F** are WT distributions from Figs 2F and 2D (with scaled Y axis in F). **G.** Spike time distribution of all spikes recorded in *shk-1* AWA under 5 s of current injection at all amplitudes. Bin size is 200 ms. The distribution in *shk-1* is not significantly different from that of WT shown in Fig 2G (right) by Kolmogorov-Smirnov Test (data not shown) despite differences in spike shape and number. **H.** Comparison of averaged action potential traces in AWA from different conditions or potassium channel mutant backgrounds. AWA spikes in WT+4AP and all *slo-1* genotypes are significantly different from WT (Table S3). Averaged *shk-1* trace is the same as in Fig 4E. Averaged WT trace (black) is the same as the red trace in Fig 2B.

In a more refined experiment, addition of the Kv1 subfamily-selective blocker 4-AP to the physiological recording solution resulted in action potentials with a larger peak amplitude, broader spike width and lower spike count (Fig 4B, H, Table S3). This observation implicates 4-AP-sensitive potassium channels in rapid membrane potential repolarization. We recorded AWA in mutants representing 11 of the 22 predicted 6TM voltage-gated potassium channel (VGKC) genes in *C. elegans* in single, double or triple mutant combinations (Fig 4C-H and Fig S3A-B), including at least one gene from each VGKC gene subfamily. The Shaker/Kv1 subfamily mutant *shk-1* had a strong phenotype resembling that of 4-AP blockade (Fig 4C, H). In *shk-1* mutants, spiking probability was increased across a range of current inputs (Fig 4E), spike amplitude and width were increased (Fig 4D, F, H), and the maximum bursting frequency was reduced to 1.8 Hz (Mean frequency: 1.4 ± 0.1 Hz, n = 9), while spike time distribution relative to stimulus remained unchanged (Fig 4G) (compare Figs 4D/2B, 4E/2F, 4F/2D and 4G/2G, respectively). These results implicate SHK-1 in action potential repolarization in AWA. No significant effect on repolarization was observed in other tested potassium channel mutants and double mutants (Fig 4H and Fig S3B, Table S3).

Calcium may also contribute to the repolarization of action potentials in AWA. When extracellular Ca^2+^ was substituted with Ba^2+^, which can pass through but does not inactivate the CaV1 calcium channel, AWA action potentials remained at the peak amplitude for the entire duration of the current injection instead of terminating spontaneously (Fig S3C). These results suggest that the activation of SHK-1 potassium channel terminates the action potential together with calcium-dependent processes such as calcium channel inactivation.

### Modeling AWA action potentials based on known and predicted channels

To begin to model the channels underlying action potential dynamics in AWA, we used whole-cell voltage-clamp recordings to determine I-V relationships of its calcium and potassium currents. Eliminating most potassium currents by Cs^+^ substitution in combination with TEA and 4-AP uncovered a small depolarizing current and leak currents (Fig 5A, B). The small depolarizing current was eliminated by adding Nemadipine-A to the extracellular solution, suggesting that it arises from EGL-19 calcium channels (Fig S4A). Since EGL-19 is rapidly, albeit partially, inactivating (Jospin et al., 2002), we subtracted steady-state current from the peak current for each voltage step to estimate presumptive EGL-19 calcium currents (Fig 5C, E). The resulting I-V relationship (Fig 5D) suggested that EGL-19 was activated near a threshold around −40 mV and peaked near −10 mV, consistent with calcium currents in previous ASER recordings (Goodman et al., 1998). The activation threshold of the inferred calcium currents matches that of AWA action potentials.

**Figure 5.**
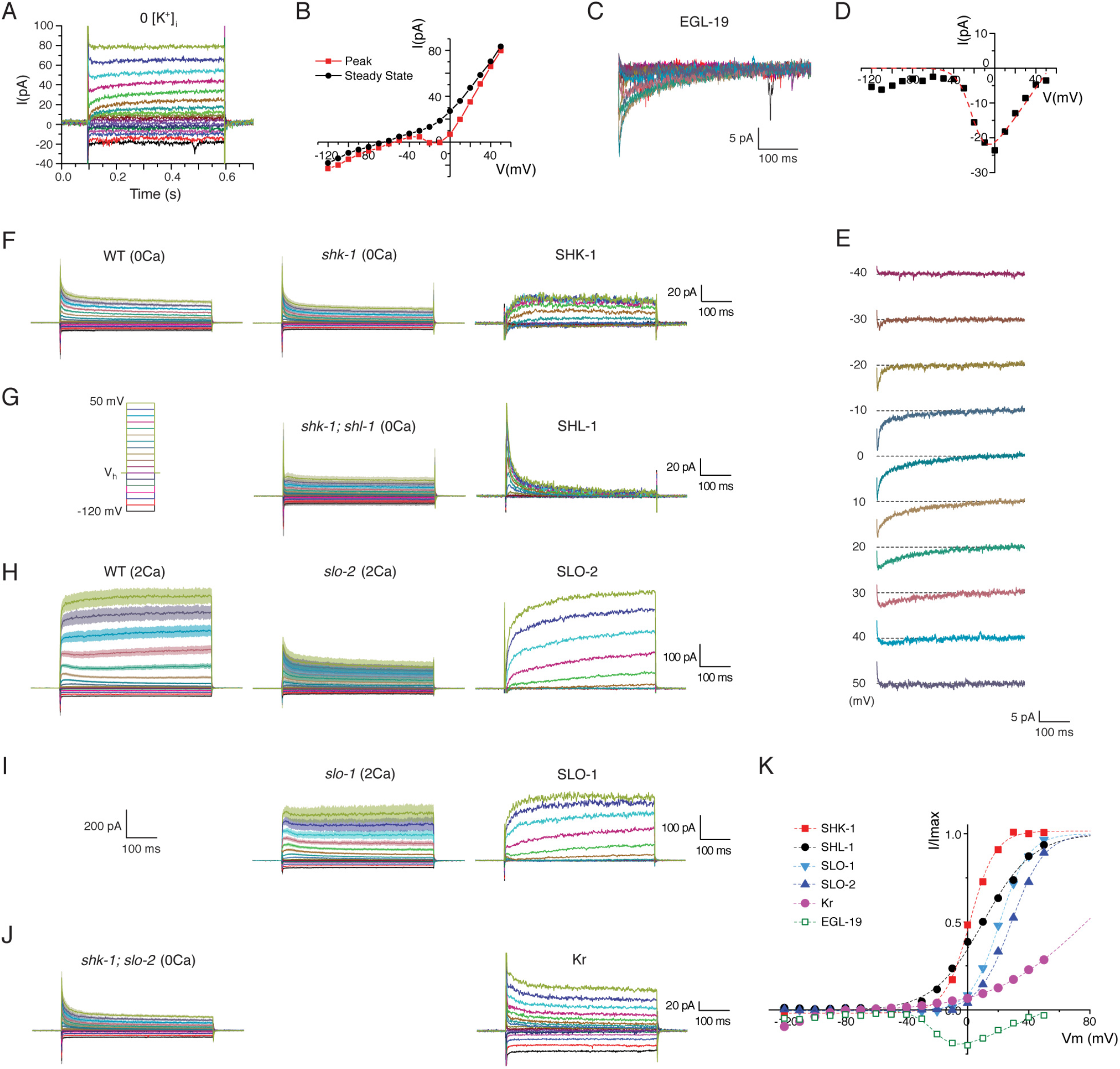
Inferred voltage-gated calcium and potassium channels underling AWA action potentials. **A.** Representative whole-cell currents in WT AWA recorded under voltage-clamp with 0 mM K^+^ intracellular solution and 150 mM TEA and 3 mM 4-AP extracellular solution. **B.** I-V relationships for peak and steady-state currents of the whole-cell currents in **A**. **C.** Superimposed EGL-19 calcium currents estimated by subtracting the steady-state currents from the peak currents in **A**. **D.** I-V relationship of estimated EGL-19 peak calcium currents and its fitting curve. **E.** Estimated calcium currents under each voltage step. **F-J.** Averaged whole-cell currents in AWA from WT and various potassium channel mutants recorded with different [Ca^2+^]_o_ (Left and middle panels) and estimated potassium currents mediated through specific potassium channels (right panels). **F.** SHK-1 currents were estimated by subtracting *shk-1* from WT in 0 mM [Ca^2+^]_o_. **G.** SHL-1 currents were estimated by subtracting *shk-1; shl-1* from *shk-1* in 0 mM [Ca^2+^]_o_. **H-I.** SLO-1 or SLO-2 currents were estimated by subtracting *slo-1* or *slo-2* from WT in 2 mM [Ca^2+^]_o_. **J.** Residual K currents (Kr) were inferred by subtracting isolated SHL-1 currents from *shk-1; slo-2* in 0 mM [Ca^2+^]_o_. Note difference in Y axis in panels **F**, **G**, and **J** (20 pA) compared to **H** and **I** (100 pA). **K.** I-V relationships of isolated currents for SHK-1, SHL-1, SLO-1, SLO-2, Kr and EGL-19, normalized to the maximum currents. Dotted lines are fitted with Boltzmann function.

The large potassium currents recorded in physiological solutions (Fig 1B, 5H) became much smaller after extracellular Ca^2+^ was removed (Fig 5F), suggesting that a subset of AWA potassium currents is calcium dependent. SHK-1 (Kv1) and SHL-1 (Kv4) currents are typically calcium-independent, and therefore should be preserved at 0 mM [Ca^2+^]_o_. We estimated the properties of potassium channels by comparing AWA in wild type to channel mutants, recognizing that this is an approximation that overlooks homeostatic effects. Subtracting the residual current in *shk-1* mutants from the wild-type whole-cell current at 0 mM [Ca^2+^]_o_ yielded a non-inactivating current inferred to be the SHK-1 current (Fig 5F), with properties consistent with previous studies (Fawcett et al., 2006; Liu et al., 2011). Subtracting the whole-cell current in *shk-1; shl-1* from that of *shk-1* at 0 mM [Ca^2+^]_o_ yielded a transient fast activating current, inferred to be SHL-1 (Fig 5G) (Fawcett et al., 2006; Santi et al., 2003).

Turning to potential calcium-dependent channels, SLO-2 is a large conductance potassium channel (Yuan et al., 2000) that is activated by Cl^-^ and Ca^2+^ entry (Liu et al., 2014). *slo-2* mutants had dramatically smaller potassium currents than wild-type at 2 mM [Ca^2+^]_o_ (Fig 5H), suggesting that SLO-2 is a dominant calcium-dependent potassium channel in AWA. Subtracting the current in *slo-2* mutants from wild-type current yielded a large slow-activating non-inactivating current inferred to be SLO-2 (Fig 5H), consistent with previous reports (Liu et al., 2011; Santi et al., 2003). SLO-1 is another large conductance calcium-activated potassium channel that plays an important regulatory role in synaptic transmission (Wang et al., 2001). *slo-1* mutants also had smaller potassium currents than wild-type at 2 mM [Ca^2+^]_o_ (Fig 5I). The SLO-1 currents inferred by subtracting the current in *slo-1* mutants from wild-type current were similar to SLO-2 currents (Fig 5I).

The I-V relationships of the four inferred potassium currents (Fig 5K) suggest that the activation voltage of SHK-1 falls narrowly above that of EGL-19, supporting the model that SHK-1 repolarizes the action potential. SHL-1 would be activated at the same voltage, but this transient current appears not to substantially influence spiking. SLO-2 and SLO-1 generate larger currents, but are activated only above 0 mV, which exceeds the effective spiking range.

After SHK-1, SHL-1, and SLO-2 currents were all subtracted from the wild-type whole cell currents, a residual fast-activating slow-inactivating potassium current remained (Fig 5J, “Kr”). The inferred Kr channel(s) were not affected by mutants for the eleven tested potassium channels, so the molecular identity of Kr is unknown. However, its properties are a central element of the AWA spiking model described below, as Kr accounts for the slow rise in membrane potential that precedes and sets the timing of the first spike (Fig 6A).

**Figure 6.**
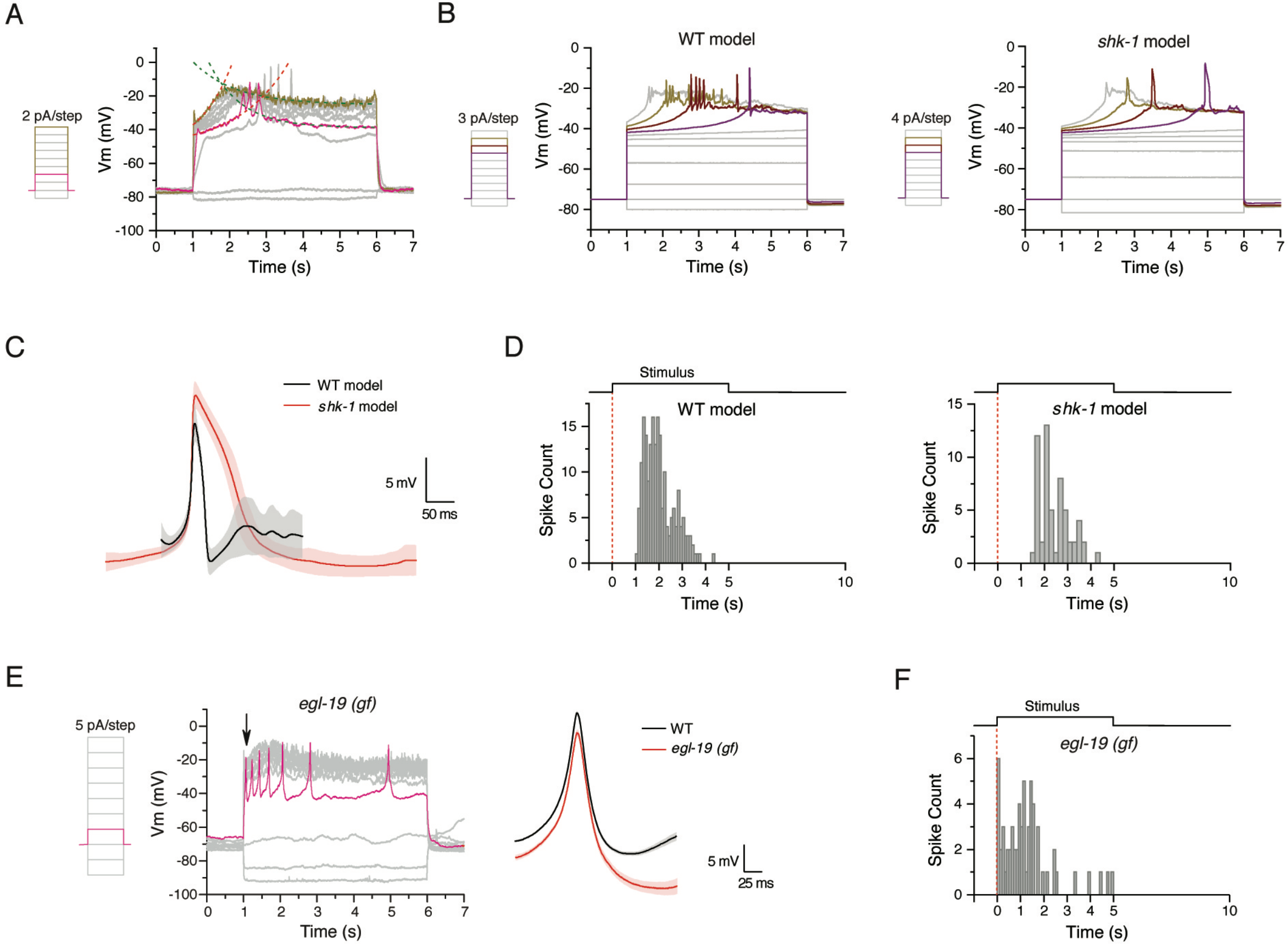
Modeling AWA action potentials based on known and unidentified channels. **A.** WT membrane potential traces (same trace as Fig 2A) showing the time dependent dynamics when action potentials fired (magenta trace) or did not fire (green trace). The voltage-dependent time course of the depolarization following the initial partial spike was well fitted with an exponential decay function (red dotted fitting curves), indicating a fast-activating and slow-inactivating K channel (Kr). The slow repolarizations (green dotted fitting curves) following the blunt peak were also well fitted with an exponential decay function and were not seen in 0 mM [Ca^2+^]_o_ (Fig 3C), suggesting that they are likely due to calcium-dependent K currents. **B-D.** Simulation of AWA action potential spikes with a Hodgkin– Huxley-type model using parameters of isolated calcium and potassium channels measured through voltage-clamp recordings. **B.** Representative action potential simulations under series of current injection steps for WT (left) and *shk-1* (right). **C.** Averaged action potentials for WT (black) and *shk-1* (red) from simulations. **D.** Spike time distribution for WT (left, n = 199) and *shk-1* (right, n = 58) from simulations. **E.** Representative action potential traces recorded from AWA in *egl-19(ad695gf)* gain-of-function mutant strain. Arrow indicates the absence of the first-spike delay. Insert: Comparison of averaged AWA action potential spikes from WT (same as the red trace in Fig 2B) and *egl-19(ad695)* (n = 24). **F**. Spike time distribution of all spikes recorded in *egl-19(ad695)* AWA under 5 s of current injection, also showing the lack of first-spike delay.

To ask whether these channels are sufficient to generate AWA action potentials, we developed a Hodgkin-Huxley type model of the AWA membrane potential (details in Experimental Procedures). The initial model incorporated the inferred properties of EGL-19, SHK-1, and SLO-1/SLO-2 channels, an additional leak current (inferred from the steady-state current in Fig 1C), and an inwardly-rectifying potassium channel (Hibino et al., 2010) that set the resting potential at −75 mV. This model gave rise to tonic spiking. Following a calculation of the bifurcation diagram that defined the spiking regime, we added the inferred slowly-inactivating Kr potassium channel that determined the delay to spiking, and a second slowly-activating Kr_a_ potassium channel to terminate spiking (Fig 6A and Experimental Procedures). To include effects of noise, both EGL-19 and Kr/Kr_a_ channels were modeled as stochastic, while other components were simulated with deterministic properties for convenience. The model showed good agreement with the measured shapes of AWA action potentials in wild-type animals, and additionally predicted the shape of action potentials in *shk-1* mutants (Fig 6B, C, compare Fig 4C, H). The model also predicted the timing and length of action potential bursts (Fig 6D, compare Fig 2G, 4G) and AWA dynamics in response to current ramps and sinusoids (Fig S4D, E, compare Fig 8A, B).

According to the model and the I-V relationships in Fig 5K, the EGL-19 calcium conductance competes with several potassium conductances to dictate the membrane potential dynamics that permit spiking. We predicted that either the removal of the Kr currents or a left shift of the activation voltage of EGL-19 would eliminate the delay to the first spike. Indeed, *egl-19(ad695)*, a gain of function mutant in which the EGL-19 I-V curve is shifted by 6 mV toward negative potentials (Laine et al., 2014), eliminated the delay to the first spike in almost all AWA recordings (Fig 6E, F). This effect supports the identification of EGL-19 as a molecular mediator of the AWA action potential upstroke.

### Natural odor stimuli may elicit AWA action potentials

AWA detects attractive volatile odors (Bargmann et al., 1993), which elicit a rise in calcium indicative of membrane depolarization (Larsch et al., 2015; Larsch et al., 2013; Shinkai et al., 2011). Exhaustive attempts to deliver odors to the dissected electrophysiological preparation were unsuccessful, so we took an indirect approach to connect action potential spikes and odor stimulation using calcium dynamics as an intermediate. In the dissected preparation, we conducted simultaneous voltage-clamp recording and calcium imaging of AWA using the genetically-encoded calcium indicator GCaMP6f. Depolarizing AWA to the spike threshold (−40 mV) resulted in smoothly increasing GCaMP6f fluorescence after a delay of approximately one second (Fig 7A); repolarizing AWA toward the resting potential resulted in fluorescence decreases without an appreciable delay. In traces that included action potentials, individual spikes were associated with an immediate increase in the slope of GCaMP6f fluorescence (Fig 7B). These results indicate that action potentials induce a sharper increase in AWA calcium than depolarization that does not drive spikes.

**Figure 7.**
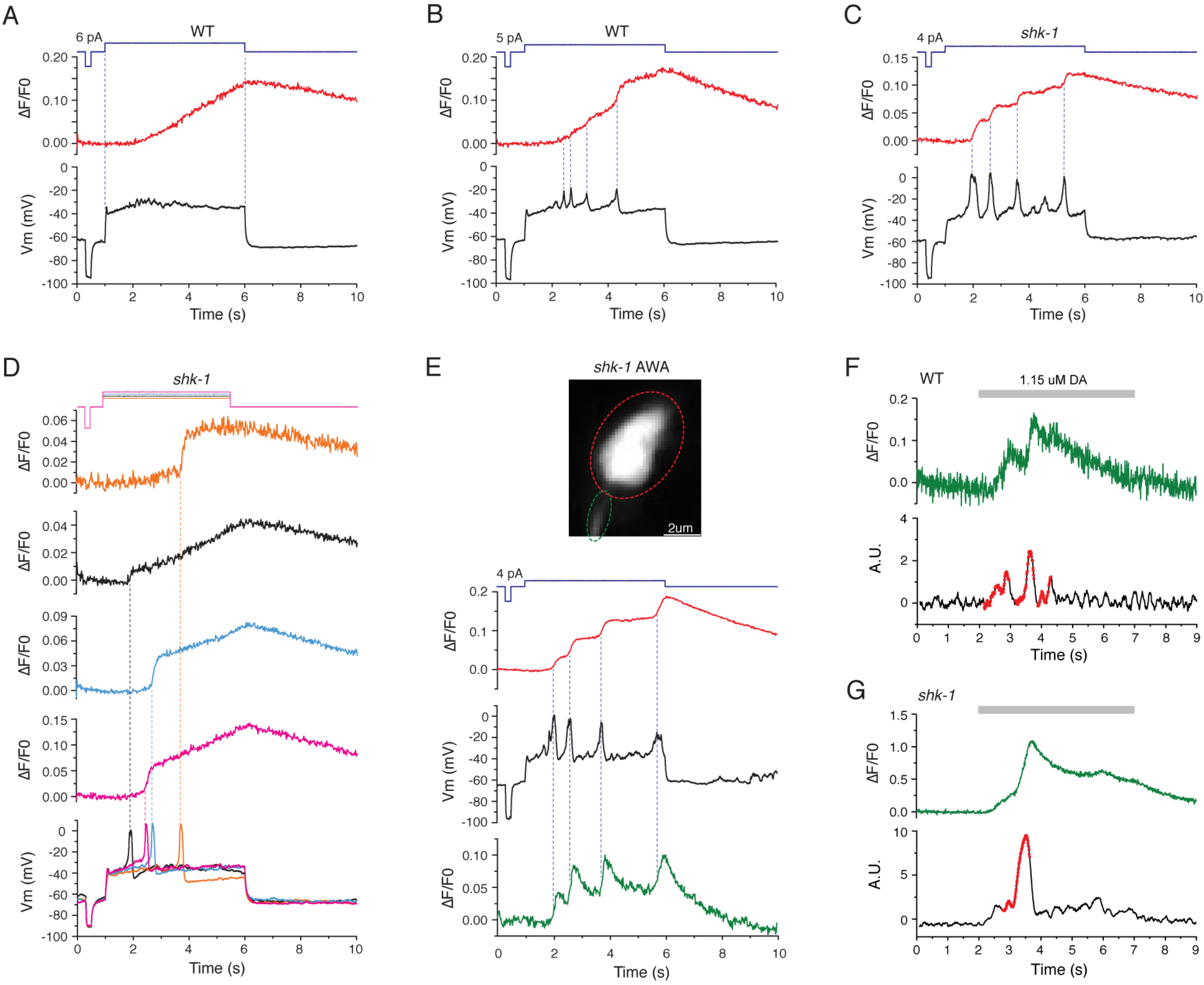
Natural odor stimuli may elicit AWA action potentials. **A-B.** Examples of simultaneous membrane potential recording (lower traces) and calcium imaging (upper traces) in AWA. **A.** When no action potentials were induced, GCaMP6f fluorescence increases linearly. Dotted lines mark beginning and end of the membrane potential change. **B.** When action potentials were induced, individual spikes were associated with an immediate increase in the slope of GCaMP6f fluorescence (marked by dotted lines). **C-D.** Example of simultaneous membrane potential recording and calcium imaging in *shk-1* AWA. Transient increase in GCaMP6f slope were correlated with individual spikes in either a recording trace with multiple spikes induced by a 5 s stimulus (**C**) or single spikes induced by four consecutive 5 s stimuli (Current injection amplitude: 4 pA (orange), 5 pA (black), 6 pA (blue) and 7 pA (pink)) (**D**). **E.** Representative simultaneous current-clamp recording (black) and calcium imaging in both the cell body (red) and axon (green) in *shk-1* AWA induced by a 5 s stimulus at 4 pA amplitude. Dotted lines indicate the correlation of each action potential with changes in the slope of calcium signals in cell body and axon. Insert: a frame of the raw video of the calcium imaging used for analysis. Red circle labels AWA cell body and green circle labels the axon. **F.** GCaMP6f fluorescence signal in a WT AWA axon evoked by a 5 s pulse of 1.15 µM diacetyl (DA), showing changes in the slope of calcium signals (upper panel). Filtering the raw trace with parameters gained from simultaneous voltage recording and calcium imaging in WT AWA generates spike-like signals (lower panel). **G.** GCaMP6f fluorescence signal in a *shk-1* AWA axon evoked by 5 s pulse of 1.15 µm diacetyl, showing a large, singular spike-like signal (lower panel). Red dotted traces in **F** and **G** were identified with the spike detection algorithm for identifying action potential spikes in WT (Fig S6) and *shk-1* (Fig S7) AWAs, respectively.

**Figure 8.**
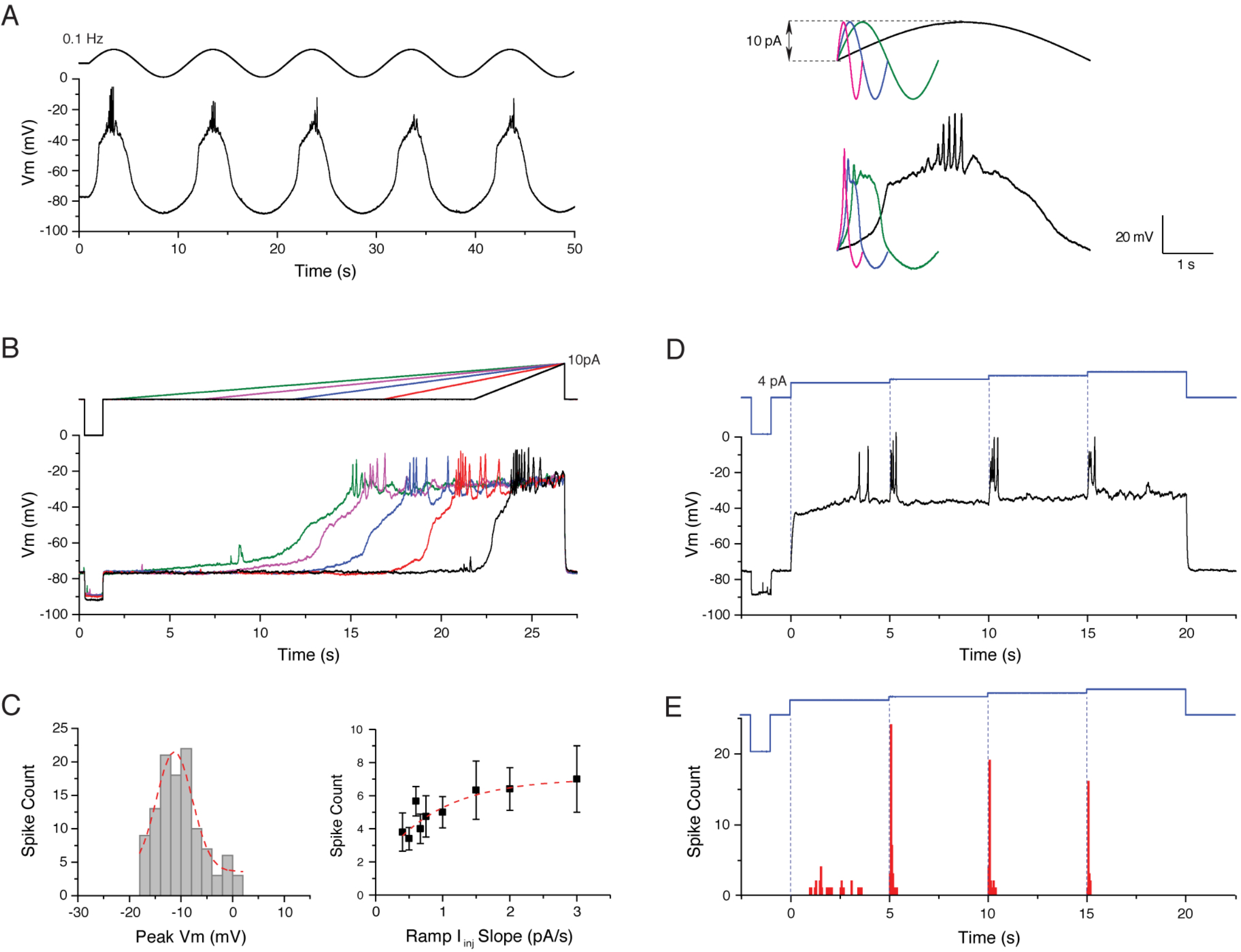
AWA action potentials may encode specific stimulus features. **A.** Action potential traces induced by 0.1 Hz sine wave stimuli (left panel). Right panel insert: close-ups of the membrane potential responses under the first cycle of sinusoidal stimuli at four different frequencies (Magenta: 2 Hz; blue: 1 Hz; green: 0.5 Hz; black: 0.1 Hz). Spikes were only fired under 0.1 Hz stimuli. **B.** Action potentials evoked by current injection ramps at different slopes in WT AWA (black: 10 pA/5 s; red: 10 pA/10 s; blue: 10 pA/15 s; pink: 10 pA/20 s; green: 10 pA/25 s). Note progressively increasing spike numbers under steeper ramp slopes. **C.** Left panel: amplitude histogram of all action potential spikes evoked by ramp stimuli at all slopes in WT AWA. Bin size is 2 mV. Red dotted line is fitted with a Gaussian distribution function (Mean = −11.3 mV, SEM = 0.4 mV, R-Square = 0.90). Right panel: correlation of spike number with the slopes of ramp stimuli. Red dotted line is fitted with an exponential decay function (R-Square = 0.64). **D.** Action potential traces induced by a 4-step current injection protocol in WT AWA. Each step is 5 s in duration. The first 4-pA step induced action potential spikes with a variable delay, while each subsequent 1 pA step induced a short burst of spikes concurrent with onset (blue dotted lines). **E.** Time distribution of the first spike in WT AWA induced by each upstep under the 4-step current injection protocol. Bin size is 50 ms, n = 37.

*shk-1* mutants have prolonged action potentials, presumably associated with prolonged EGL-19 channel opening. In *shk-1* mutants, the transient increase in GCaMP6f slope was enhanced both for multiple spikes (Fig 7C) and for single spikes (Fig 7D). This signal was further enhanced when measuring calcium in the AWA axon, where calcium clearance is accelerated compared to the soma (Fig 7E).

Based on the GCaMP6f signals induced by smooth depolarization (Fig 7A) versus spiking (Fig 7B-E), we sought evidence that natural odor stimuli could induce AWA action potentials in intact animals. The bacterial metabolite diacetyl is a potent stimulator of AWA calcium transients and AWA-driven chemotaxis behavior (Bargmann et al., 1993). Applying derivative and amplitude thresholds and deconvolution to diacetyl-induced GCaMP6f signals supported the presence of action potentials in both wild-type (Fig 7F) and *shk-1* (Fig 7G) animals. In agreement with the predictions from electrophysiology, the inferred action potentials were consistently smaller and more frequent in wild-type than in *shk-1* animals. These results suggest that diacetyl odors can elicit AWA action potentials in intact animals.

### AWA action potentials may encode specific stimulus features

Stimuli in natural environments can fluctuate on different temporal and spatial scales. To identify conditions that best elicit AWA action potentials, we conducted electrophysiological recordings while modulating current in different patterns.

Among a variety of ramps and modulated stimuli, we found that sine wave stimuli from −10 to +10 pA at 0.1 Hz generated AWA action potential bursts reliably across trials (Fig 8A). By contrast, 0.5, 1, or 2 Hz sine wave stimuli – chosen to match the timescale of *C. elegans* sinusoidal movement – were ineffective at inducing action potentials (Fig 8A and Fig S5A). To ask whether the rate of stimulus change is a key mediator of AWA spiking, we challenged AWA with current injection ramps at slopes ranging from 0.4 to 3 pA/s (Fig 8B). In both wild-type and *shk-1* animals, spikes were readily induced by these slow ramps (Fig 8B and Fig S5B), but not by a fast ramp at 25 pA/s (Fig S5C). At slow ramps, the number of spikes had a modest positive correlation with the slope of the stimulation, but spike amplitude was unchanged (Fig 8C). Together, these results suggest that slowly rising stimuli result in reliable AWA spiking.

Calcium imaging studies have suggested that AWA preferentially detects spaced upsteps in odor concentrations, rather than absolute odor thresholds (Larsch et al., 2015). To ask whether this property is reflected in action potentials, we elicited a burst of action potentials with a 4 or 5 pA current, and then delivered additional 1 or 2 pA-increments of current at 5 second intervals (Fig 8D). Once spiking had been induced, subsequent upsteps could also evoke spikes (Fig 8D). In contrast with the first burst of spikes, these upstep spikes appeared without a delay (Fig 8D, E, Fig S5D). Thus AWA can faithfully report small stimulus upsteps by firing immediate spike bursts, in a manner dependent on the stimuli history of the cell.

## Discussion

Unlike previously-characterized *C. elegans* neurons, the AWA neuron fires all-or-none calcium-based action potentials. These spikes are reliably elicited by specific current stimuli, and based on indirect calcium measurements, they appear to be elicited by natural odor stimuli as well. The action potentials are mediated by a CaV1 voltage-gated calcium channel, EGL-19, and a Kv1 voltage-gated potassium channel, SHK-1, and their properties are shifted in instructive ways by gain- and loss- of function mutations in these channels.

A delay before the first AWA spike appears to be caused by a slowly-inactivating potassium current. Testing a variety of mutants failed to identify the molecular basis of this current, which may be analogous to “A currents” that cause a first spike delay in other vertebrate and invertebrate neurons (Chu et al., 2003; Dekin and Getting, 1987; Kawaguchi, 1995; Zhao and Wu, 1997). Termination of the burst can be modeled with a slowly activating, high voltage-activated potassium channel, along with inactivation of the EGL-19 channel.

The membrane potential bistability and first-spike delay in AWA provide an elegant solution to the “small-cell limit” problem that has puzzled theorists: for small cells with high impedance, the stochastic opening of a single channel can cause voltage fluctuations that trigger an action potential, which would make firing so irregular that coding schemes based on either firing rate or spike timing are essentially unworkable (Faisal et al., 2005; Lockery and Goodman, 2009; Strassberg and Defelice, 1993). Our data indicates that membrane potential must be depolarized to the plateau state for at least 300 ms for AWA to fire action potentials. This delay far exceeds the open time of most single channels, effectively preventing transient single channel noise from triggering spikes.

### Computation and neuronal diversity

The electrophysiological properties of most *C. elegans* neurons are unknown. Our limited survey enumerated three types of non-spiking neurons and a fourth unexpected spiking neuron, suggesting that additional biophysical diversity awaits discovery in the *C. elegans* nervous system.

Among the biophysical properties in *C. elegans* neurons observed in this study, the membrane potential bistability in AFD and AWA is likely shared by multiple cell types. The flat region between −80 and −30 mV on the I-V relationship resembles the previously reported high phenomenological resistance region in AFD (Ramot et al., 2008), ASER (Goodman et al., 1998), RMD (Mellem et al., 2008), AWC (Ramot et al., 2008) and ASH (Geffeney et al., 2011). This high-resistance region in AWA is associated with a dearth of voltage-gated channel opening. Bistability is a common theme in many other neurons, including vertebrate hippocampal neurons, cortical neurons, and cerebellar Purkinje cells (Loewenstein et al., 2005). Other *C. elegans* neuronal classes described here include the outwardly rectifying AIY neurons, previously reported in Faumont et al., 2006, whose features resemble those reported for VA5 and VB6 motor neurons (Liu et al., 2017; Liu et al., 2014), and the near-linear RIM neurons, whose features resemble those reported for AVA (Lindsay et al., 2011; Liu et al., 2017; Mellem et al., 2008), PLM (O’Hagan et al., 2005) and AVE (Lindsay et al., 2011) neurons. The nearly passive behavior of RIM neurons is surprising given their bistability in calcium imaging experiments (Gordus et al., 2015), pointing to the importance of the network in conferring distinct up- and downstates on RIM.

The *C. elegans* genome encodes 74 predicted potassium channels, a remarkable repertoire for just 302 neurons. Reporter gene data and single-cell RNA sequencing data suggest a considerable diversity in their expression patterns (Cao et al., 2017; Salkoff et al., 2005). At the extreme, each neuronal cell type could express a unique channel composition with unique sensitivity and computing power to suit its function.

EGL-19 and SHK-1 are broadly expressed in *C. elegans* neurons, so it may be that neurons other than AWA can spike in stimulation regimes that remain to be identified. Indeed, the spikelets that were reported in an *sra-11-*expressing *C. elegans* interneuron resemble those described here in their amplitude and duration (Faumont et al., 2012), and unidentified neurons in the ventral nerve cord of *Ascaris* nematodes may also spike (Davis and Stretton, 1992).

### AWA spiking as a filter of odor stimuli

What information about the odor stimulus is translated into spikes? When *C. elegans* crawls across a bacterial lawn in a sinusoidal waveform, its undulatory head swings create a natural temporal oscillation in odor concentrations at its nose tip (Kato et al., 2014; Kocabas et al., 2012). We considered that this oscillation might induce phase-locked spiking in AWA, but found that the 0.5 - 2 Hz oscillation characteristic of the head swings did not trigger action potentials. Instead, the delay to the first spike may function as a low-pass filter to high frequency fluctuations, including those generated by the worm’s own head swings. Slow sustained changes in odor concentration, and the increases caused by the animal’s directed locomotion, appear more likely to induce spiking in AWA.

Neuronal firing patterns are typically modeled as either rate codes or temporal codes (Gerstner and Kistler, 2002; Sarpeshkar, 1998). The modest modulation of the spike frequency by stimulus strength (Fig 2J) and our bifurcation model indicate that AWA spiking is most consistent with class 2 excitability based on the Hodgkin classification of isolated crustacean axons (Hodgkin, 1948). Action potentials of this type fire in a narrow frequency band and are relatively insensitive to changes in the stimulus strength, incompatible with a rate-coding scheme (Izhikevich, 2007). Rate coding would also be degraded by the stochastic and self-terminating nature of action potential bursting. Instead, AWA spiking appears most consistent with a filtering function that requires a stimulus of appropriate intensity and duration. The transition between the resting state and plateau state sets the required stimulus intensity, and the first-spike delay serves as an internal time reference signal for tracking stimulus duration. Temporal coding by first-spike delay is also observed in mitral/tufted cells in the mouse olfactory bulb, albeit on a roughly ten-fold faster time scale (Margrie and Schaefer, 2003). After AWA enters the plateau state, small depolarizing inputs spaced a few seconds apart can trigger spikes without a delay. Taking both the bistability and delayed firing features into consideration, we suggest the following: when an animal encounters a suprathreshold attractive odor stimulus for the first time, AWA fires action potentials when the odor stimulus is maintained beyond the first-spike latency, allowing the animal to ignore noisy or transient stimuli. Maintaining AWA membrane potential in the plateau state induces excitability changes that terminate firing, but allow increasing odor concentrations to trigger spikes without delay, enabling odor gradient climbing behavior.

A simulation of *C. elegans* chemotaxis using an artificial neural circuit for contour tracking found that a hypothetical neural network with a pair of artificial spiking neurons as the primary gradient detector had higher chances of identifying the set-point, and better tracking efficiency, than an equivalent non-spiking network with graded sensory neurons (Santurkar and Rajendran, 2014). Whether action potentials in AWA increase chemotaxis efficiency in real animals remains to be tested by identifying mutations or conditions that selectively disrupt action potentials in AWA. More complicated possibilities can also be explored. For example, in mitral/tufted cells of the mouse olfactory bulb, first-spike latency relative to the start of a sniffing action can encode odor identity (Margrie and Schaefer, 2003). Similarly, it may be informative to ask whether different odors induce AWA action potentials with different spiking thresholds or temporal features.

The identification of spiking neurons marks a shift in our conception of information processing in *C. elegans*. Understanding the transition from single-neuron to population dynamics is the next step in assessing the significance of AWA spiking. Global brain dynamics, in which groups of neurons exhibit coordinated activities, are associated with behavioral states in *C. elegans* (Kato et al., 2015). Future experiments should reveal the roles that spiking neurons play in dynamic network activity and its global state transitions. Do a small number of spiking neurons contribute to a central pattern generator, or does synchronized spiking enable dynamic switching at a global scale?

Digital compression of information with action potentials is inherently lossy, while analog multiplexing of information can expand one neuron into many computational units. With a small number of neurons, *C. elegans* is best served by heavily reliance on analog coding. For example, the local compartmentalized coding in the RIA integrating neuron allows one neuron to carry both sensory and motor streams of information (Hendricks et al., 2012). Understanding the functions of AWA spikes provides a sharply focused example in which to ask where action potentials are most critical in neural coding.

## Author Contributions

Q.L. and C.I.B. designed experiments and oversaw the project. Q.L. designed and conducted electrophysiological experiments. P.B.K. designed and wrote the spike detection and analysis algorithm and conducted computational modeling. M.D. designed and conducted calcium imaging experiments. Q.L., P.B.K., M.D. and C.I.B. analyzed and interpreted data. Q.L. and C.I.B. wrote the paper.

## Acknowledgments

We thank Andrew Gordus, Larry Abbott, Jim Hudspeth, and many lab members for scientific discussions, Bob Horvitz, Zhao-Wen Wang, Shawn Lockery, Miriam Goodman, and Piali Sengupta for comments on the manuscript, and the *Caenorhabditis* Genetics Center (CGC), Zhao-Wen Wang, Larry Salkoff, Michael Nonet and Erik Jorgensen for strains. This work was supported by a Kavli NSI Pilot Grant to Q.L, the Howard Hughes Medical Institute, of which C.I.B. was an investigator, and the Chan Zuckerberg Initiative.

## Experimental Procedures

### Nematode growth

All strains were maintained at room temperature (22-23°C) on nematode growth medium (NGM) plates, seeded with *E. coli* OP50 bacteria as a food source (Brenner, 1974). Wild-type animals were the Bristol strain N2. A complete strain list is provided in Table S1.

### Electrophysiology

Electrophysiological recording was performed largely as previously described (Goodman et al., 1998; Liu et al., 2009). Briefly, an adult animal was immobilized on a Sylgard-coated (Sylgard 184, Dow Corning) glass coverslip in a small drop of DPBS (D8537; Sigma) by applying a cyanoacrylate adhesive (Vetbond tissue adhesive; 3M) along the dorsal side of the head region. A puncture in the cuticle away from the head was made to relieve hydrostatic pressure. A small longitudinal incision was then made using a diamond dissecting blade (Type M-DL 72029-L; EMS) between two pharyngeal bulbs along the glue line. The cuticle flap was folded back and glued to the coverslip with GLUture Topical Adhesive (Abbott Laboratories), exposing the nerve ring. The coverslip with the dissected preparation was then placed in a custom-made open recording chamber (~1.5 mL volume) and treated with 1 mg/mL collagenase (type IV; Sigma) for ~10s by hand pipetting. The recording chamber was subsequently perfused with desired extracellular solution with a custom-made gravity-feed perfusion system for ~10 mL. All experiments were performed with the bath at room temperature. An upright microscope (Axio Examiner; Carl Zeiss, Inc.) equipped with a 40 X water immersion lens and 16 X eyepieces was used for viewing the preparation. Neurons of interest were identified by fluorescent markers. Borosilicate glass pipettes (BF100-58-10; Sutter Instruments) with resistance (RE) = 10–15 MΩ pre-pulled using a laser pipette puller (P-2000; Sutter Instruments) were back-filled with desired intracellular solution to use as electrodes. A motorized micromanipulator (PatchStar Micromanipulator; Scientifica) was used to control the electrodes. Whole-cell current clamp or voltage clamp experiments were conducted on an EPC-10 amplifier (EPC-10 USB; Heka,) using PatchMaster software (Heka). Two-stage capacitive compensation was optimized at rest, and series resistance was compensated to 50%. Analog data were filtered at 2 kHz and digitized at 10 kHz or 50 kHz. As the quality control, only those patch-clamps with seal resistance above 1 GOhm and uncompensated series resistance below 100 MOhm were accepted for further analysis (seal resistance ranged from 1 to 9 GOhm and series resistance ranged from 20 to 60 MOhm for most accepted recordings). To ensure reliable spiking induction, a short −10 pA hyperpolarizing pre-pulse was added to each current injection step in the stimulation protocol in many experiments (e.g. Fig S1C).

### Recording solutions

The standard pipette solution was (all concentrations in mM) [K-gluconate 115; KCl 15; KOH 10; MgCl_2_ 5; CaCl_2_ 0.1; Na_2_ATP 5; NaGTP 0.5; Na_2_cGMP 0.5; cAMP 0.5; BAPTA 1; Hepes 10; Sucrose 50] (pH was adjusted with KOH to 7.2, osmolarity 320-330 mOsm) and the standard extracellular solution was [NaCl 140; NaOH 5; KCl 5; CaCl_2_ 2; MgCl_2_ 5; Sucrose 15; Hepes 15; Dextrose 25] (pH was adjusted with NaOH to 7.3, osmolarity 330-340 mOsm). For recording calcium currents in 0 mM [K^+^]_i_ condition (Fig 4A and Fig 5A), pipette solution was [CsCl 140; TEA-Cl 5; TES 5; CaCl_2_ 0.1; Na_2_ATP 5; NaGTP 0.5; Na_2_cGMP 0.5; cAMP 0.5; BAPTA 1; Hepes 10; Sucrose 30] (pH was adjusted with TEA-OH to 7.2, osmolarity 320-330 mOsm) and the extracellular solution was [TEA-Cl 150; CaCl_2_ 5; 4AP 3; Sucrose 10; Hepes 15; Dextrose 25] (pH was adjusted with TEA-OH to 7.3, osmolarity 330-340 mOsm). For recordings in 1, 5 and 8 mM [Ca^2+^]_o_ (Fig 3D and S2A), extracellular CaCl_2_ was adjusted accordingly. For recordings in 0 mM [Ca^2+^]_o_ (Fig 3C), extracellular CaCl_2_ was replaced with 2 mM EGTA. For recordings in 2 mM [Ba^2+^]_o_ (Fig S3C), extracellular CaCl_2_ was replaced with 2 mM BaCl_2_. For recordings in 0 mM [Na^+^]_o_ (Fig 3B), extracellular NaCl was replaced with 140 mM NMDG. Liquid junction potentials were calculated and corrected before recording for each pair of solutions used. For Nemadipine-A application, Nemadipine-A (Sigma) was first dissolved in DMSO then further diluted in extracellular solutions to the working concentration (contained 0.04% DMSO at 10 µM Nemadipine-A).

### Simultaneous electrophysiology and calcium imaging

The electrophysiology recording setup was as described above. GCaMP fluorescence was captured with a CoolSNAP HQ2 Camera (Photometrics) controlled by MetaMorph software. The onset of imaging acquisition and electrophysiology recording was synchronized by a TTL signal sent through Patchmaster. Illumination was a SpectraX Lumencor solid-state light source. Patch-clamping was performed under DIC illumination, and the sample was switched to GFP illumination once a whole-cell recording configuration was formed to start simultaneous recording. Imaging was set at 20 MHz, binning at 2 and 50 fps. Tiff imaging stacks were analyzed using MetaMorph by drawing circles around the areas of interest (cell body or axon segment) and measuring average intensity for each frame. Time sequence of membrane voltage and fluorescence intensity measurements were then aligned by time 0 and plotted (Fig 7A-E).

### Spike detection and analysis

Action potential spikes for averaging and measurement were identified using a customized spike-detection algorithm written in Python. For the calculation of the average upstroke, downstroke, spike shape, spike timing distributions, and other statistics, action potentials were identified automatically in the data using a moving window and two threshold criteria for the rising and falling slope of the potential. The thresholds and the width of the window were tuned by hand on representative data, shown in Fig S6 and S7.

A burst was defined as two or more spikes fired within 2 s. Spiking frequency was calculated by taking the reciprocal of the averaged spike intervals for each burst. Identified spikes were aligned by the maximum slope of the upstrokes for averaging. False positives resulting from obvious noise and misalignment were excluded by visual inspection. WT action potentials with adjacent spikes within 150 ms prior or after the peak were also manually excluded for minimizing the baseline and AHP distortion. The averaged trace of 149 “good spikes” shown in Fig 2B was used for comparison with other genotypes. 19 well-separated action potential spikes were separately analyzed as “single spikes” for Fig 2C. Since most action potentials in *shk-1* are well-separated from one another, no distinctions were made between single spikes and spikes in bursts. All verified spikes were used for analysis in Fig 4D.

### Statistics and graphing

Statistics, curve fitting and graphing were conducted using OriginPro (OriginLab). I-V relationships were measured using Fitmaster (HEKA) and exported to OriginPro. Electrophysiological recording traces shown in all figures were re-sampled at 2.5kHz by averaging adjacent data points for reducing file size.

Curve fitting equations:

Gaussian function (Fig 2D, 2E, 2F, 2I, 4E, 4F, 8C):

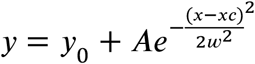

Exponential decay function (Fig 2H, 2J, 6A, 8C):

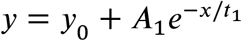

Boltzmann function (Fig 5K):

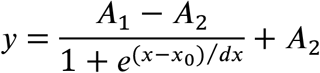

Calcium I-V curve fit function (Fig 5D):

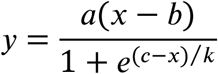

Current subtractions (Fig 5F-J) were performed after normalizing the whole-cell currents to the maximum hyperpolarizing currents (assuming that hyperpolarizing currents are not affected by the K channels of interest).

### Odor-evoked calcium imaging and data analysis

Odor-evoked calcium imaging experiments were performed on young adult N2 or *shk-1(ok1581)* animals. GCaMP6f, chosen for its fast calcium dissociation kinetics (Chen et al., 2013), was codon-optimized and expressed in the AWA neuron under the *gpa-6* promoter (pMD3 = gpa-6P::GCaMP6f). Calcium dynamics of the AWA cell body and axon were imaged using single-animal microfluidics recording chambers (Chronis et al., 2007) connected to reservoirs of S Basal buffer and freshly prepared 1.15 µM diacetyl (Sigma D3634) in S Basal. Switching between buffer and diacetyl was achieved using a solenoid valve controlled by Metamorph software, as previously described (Chronis et al., 2007). Well-fed animals were paralyzed with 1 mM (-)-tetramisole hydrochloride (Sigma L9756) for 10 minutes before being loaded into the chamber. Two 5 second diacetyl pulses, separated by 5 minutes, were recorded per animal for a total of 40 N2 videos and 56 *shk-1* videos. Videos were recorded at 100 fps under continuous 484-492 nm light through a 40X 1.3 NA Zeiss Plan-APOCHROMAT objective on a Zeiss Axiovert 100TV microscope. An Andor iXon+ DU-987 EMCCD camera and Metamorph software were used for image acquisition. The fluorescence values were extracted from each TIFF stack using ImageJ software. The mean value of 1 s baseline fluorescence before the onset of diacetyl pulse was used as F0. ΔF was calculated as F-F0 and all imaging graphs were plotted as ΔF/F0.

### Filtering and spike identification

To identify action potentials in AWA GCaMP time courses, we deconvolved the fluorescence data by a decaying exponential kernel with a 1 second time constant and smoothed the result by averaging over a 200 millisecond moving window. Spikes were then identified using the method described above, with thresholds tuned to give correct results on the ground truth data when GCaMP fluorescence and membrane potential were measured simultaneously (see Fig 7). The data was normalized to a scale of zero to one for spike detection, but for plotting the original fluorescence scale is restored. The units represent a change in fluorescence intensity above baseline, and show that the detected spikes in *shk-1* are larger than in wild type.

### Modeling and simulation of AWA spikes

The dynamics of the membrane potential, *ν*, are governed by an equation of the form

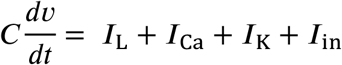

where *C* is the membrane capacitance, and the terms on the right represent leak current, calcium current, potassium current, and input current, respectively. The leak current is given by Ohm’s law: ***I***_L_ = g_L_(*ν-ν_L_*). We began by modeling the SHK-1 potassium channel and the EGL-19 calcium channel. The SHK-1 potassium current is given by

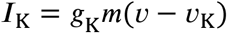

where g_K_ is the overall conductance (fit from the voltage clamp data in Fig 5F),*ν*_K_ is the potassium reversal potential (set to the usual value of −84 mV), and *m* is a gating variable whose dynamics are described by the equations

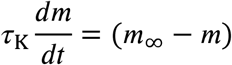

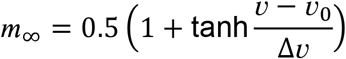

where τ_K_ is a characteristic time scale fit from the voltage clamp dynamics in Fig 5F, and the steady-state function *m*_∞_ is made to match the red curve in Fig 5K by fitting the threshold voltage *ν*_0_ and slope Δ*ν*

The EGL-19 calcium channel was modeled in a similar fashion. The steady-state function of the gating variable was represented as a product of two sigmoid functions of the form *m_∞_* to reflect the opening and subsequent closing of the channel as membrane potential increases. A simulation of these two channels alone with a realistic leak current (estimated from the IV curve in Fig 1C) and all other parameters fit directly from the voltage clamp data in Fig 5 shows spontaneously generated action potential spikes, lending credence to the hypothesis that EGL-19 and SHK-1 are the primary channels responsible for shaping the action potentials in AWA. To fit the high inward current evident in the IV curve at higher membrane potentials (Fig 1C), we added two channels with threshold voltages and slopes similar to those measured for the SLO-1/SLO-2 channels, but the model required a higher conductance than was measured, suggesting that additional potassium channels open at depolarized membrane potentials. The threshold voltages and slopes were modified slightly to give a better fit to the whole cell I-V curve (Fig 1C). Finally, we added an inwardly-rectifying potassium channel with a conductance tuned to match the resting potential in AWA (about −75 mV). This model predicts tonic spiking in a small range of input currents from about 12 to 16 pA, only slightly larger than the values of current input that elicit bursts of spikes in AWA.

We also modeled self-inactivation of the calcium channels, as observed in the voltage clamp data taken in the presence of K-channel inhibitors (Fig 5B). This was accomplished by allowing the calcium channel to have two states, with the second state having a lower conductance than the first by a factor *f*. The correct conductances were estimated by fitting a sigmoidal function plus a linear leak to the I-V curves in Fig 5B and taking the difference. The time scale of inactivation was voltage-dependent and fit by matching the relaxation times of the curves in Fig 5A.

To model the bursting structure of action potentials in AWA, we added two potassium channels, one with a slowly inactivating current (which we call Kr), and one with a slowly activating current (which we call Kr_a_). These channels together represent the slowly modulated Kr currents that gate the initiation and termination of spiking. To determine the properties of these channels, we first calculated the bifurcation diagram of the model without the Kr and Kr_a_ current, and then added an unknown potassium conductance in the form of an additional term of the form g_K_(*ν-ν_K_*). We used numerical continuation in the PyDSTool package (Clewley, 2012) to find the line of Hopf bifurcations in the *I*-*g*_K_ plane. The results (Fig S4B) show that spiking is possible in a bounded tongue-shaped region. We tuned the threshold and conductance of the Kr channel such that, for realistic values of the input current, the slow inactivation of Kr caused *g*_K_ to drop into the spiking region after about 1 second, initiating a burst. The corresponding parameters for the Kr_a_ channel were tuned so that the slow activation of Kr_a_ caused *g*_K_ to rise out of the spiking region, terminating the burst after a realistic number of spikes (2-5, see Fig 6B). To match the length of bursting at different input currents, we allowed the activation time of the Kr_a_ channel to depend on voltage, such that activation occurred more quickly at higher potentials. The path of potassium conductance values at different input currents (Fig S4C) was also designed to give reasonable spiking amplitudes across the range of allowed inputs (see Fig S4B). The earlier onset time and slightly higher frequency of bursting arise naturally from these elements. The potassium conductances in Fig S4C are computed from a fully deterministic equation, and they actually fall outside the allowed spiking region. This occurs because the addition of noise blurs out the bifurcation boundary, so that spiking will occur outside of the region where it is allowed deterministically.

We accounted for noise using a hybrid algorithm in which opening of the calcium and Kr channels was stochastic, while other components were simulated with the deterministic equations. The opening or closing of a calcium/Kr channel was treated as a Poisson process with average rate given by the deterministic equation, and reasonable noise effects were obtained for 1000 calcium channels and 500 of each Kr channel per cell. The *shk-1* mutant was simulated with 2500 of each Kr channel, due to the increased potassium conductance. A representative example is shown in Fig 6B. The model qualitatively predicted the *shk-1* mutant phenotype by setting the SHK-1 conductance to zero and increasing the conductance of the “SLO-like” and Kr channels (Fig 6C). The model also showed reasonable agreement with the measured shapes of action potentials in N2 and *shk-1* animals and the timing and length of action potentials bursts (Fig 6C-D).

There are a few notable discrepancies between the model and the data. The agreement between the spike shape in the model and in the N2 data is reasonably good, but the spike amplitude is smaller in the *shk-1* version of the model than in the *shk-1* data. The values of the input current at which spiking occurs are somewhat higher in the model than in AWA. It is likely that additional channels in AWA affect its dynamics. More interestingly, the bifurcation structure in AWA appears slightly different than in the model. In AWA, the spike amplitude typically increases over the course of a burst, in the model the opposite is true – the amplitude of spikes decreases as potassium conductance g_K_ increases. The model would therefore fit the amplitude data if a burst were initiated by an *increasing* potassium current crossing into the spiking region from below. However, this would be inconsistent with the curves shown in Fig 6A.

The model reproduces the qualitative features of AWA dynamics. In particular, the model shows delayed transient bursts of action potentials in a narrow range of input currents. We also simulated the model in the presence of linearly increasing and sinusoidally varying input currents (see experimental data in Fig 8) and obtained small bursts of spikes for low-frequency sinusoidal input and delayed bursts in response to current ramps of varying slope (simulations in Fig S4D-E).

The equations describing the calcium and potassium current are below. These together with the equations in the first paragraph of this section constitute the full model. The parameter values used are in Table S4. In cases where a parameter is changed between the N2 and *shk-1* versions of the model, both values are shown.

Calcium current:

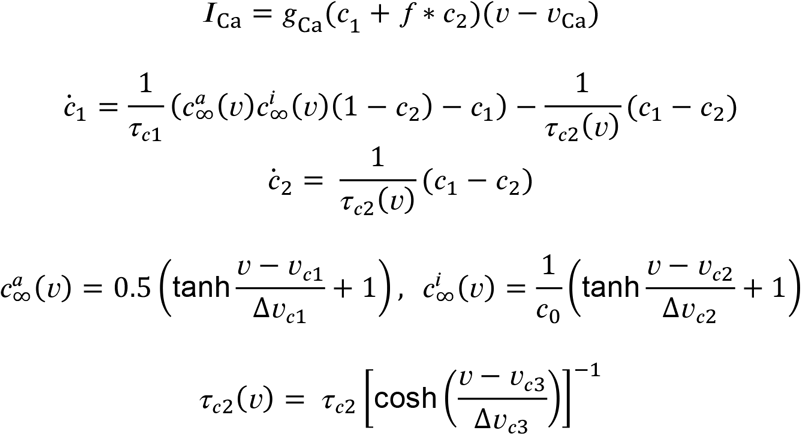

Potassium current:

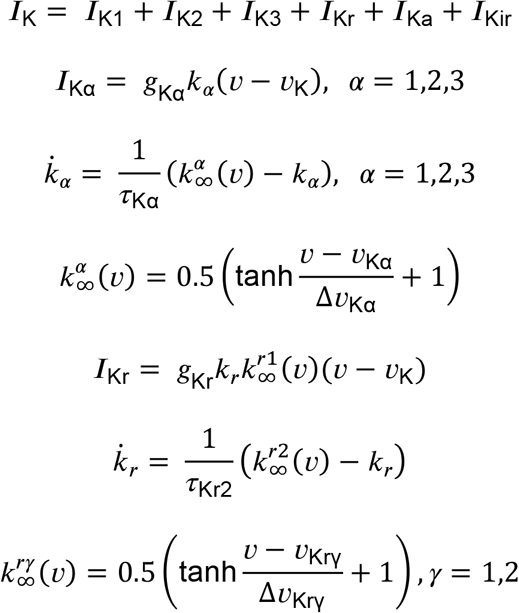

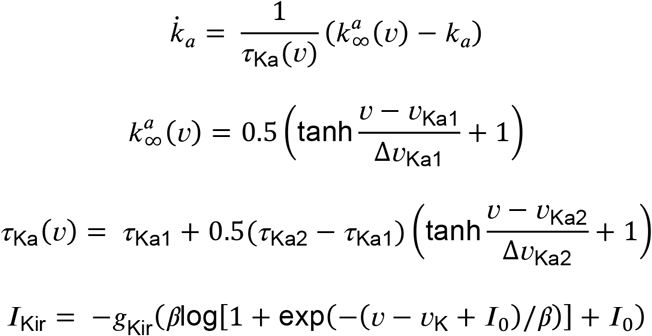

## Supplemental materials

**Table S1.** Strain list.

| Strain | Genotype | Comment | Figure |
| --- | --- | --- | --- |
| CX3262 | *kyls37[odr-10::GFP]* | *AWA::GFP* | Fig 1-6, 8, S1-S6 |
| CX12516 | *kyEx3488[gcy-13::GFP 100ng + unc-122::dsRed 10ng]* | *RIM::GFP* | Fig 1 |
| CX13495 | *kyEx4059[mod-1::GFP 50ng + unc-122::dsRed 10ng]* | *AIY::GFP* | Fig 1 |
| PY1322 | *oyls18[gcy-8::GFP]* | *AFD::GFP* | Fig 1 |
| CX17063 | *unc-13(e51); kyls37* | *unc-13; AWA::GFP* | Fig 3A |
| CX4312 | *osm-9(ky10); kyls37* | *osm-9; AWA::GFP* | Fig 3G, H |
| CX16801 | *egl-19(n582); kyls37* | *egl-19; AWA::GFP* | Fig 3E |
| CX6889 | *unc-2(e55); kyls37* | *unc-2; AWA::GFP* | Fig 3G, S2B |
| CX17062 | *cca-1(ad1650); kyls37* | *cca-1; AWA::GFP* | Fig 3G, S2B |
| CX17189 | *nca-1(gk9); nca-2(gk5); kyls37* | *nca-1; nca-2; AWA::GFP* | Fig 3G, S2B |
| CX16803 | *shk-1(ok1581); kyls37* | *shk-1; AWA::GFP* | Fig 4C-G, 4H, 5F, S5B, D, S7 |
| CX16802 | *slo-2(nf101); kyls3* | *slo-2; AWA::GFP* | Fig 4H, 5H, S3A |
| CX16955 | *shl-1(ok1168); kyls37* | *shl-1; AWA::GFP* | Fig 4H, S3A |
| CX17086 | *slo-1(eg142); kyls37* | *slo-1; AWA::GFP* | Fig 4H, 5I, S3A |
| CX17216 | *kcnl-2(ok2818); kyls37* | *kcnl-2; AWA::GFP* | Fig S3A |
| CX17187 | *egl-2(rg4); kyls37* | *egl-2; AWA::GFP* | Fig S3A |
| CX17326 | *shw-1(r1159); kyls729* | *shw-1; AWA::GFP* | Fig S3A |
| CX17289 | *egl-36(n2332sd; sa577); kyls37* | *shw-2; AWA::GFP* | Fig S3A |
| CX17087 | *exp-3(n2372); kyls37* | *exp-3(gf); AWA::GFP* | Fig S3A |
| CX17039 | *slo-2(nf101); shk-1(ok1581); kyls37* | *slo-2; shk-1; AWA::GFP* | Fig 4H, 5J, S3B |
| CX17085 | *shl-1(ok1168); shk-1(ok1581); kyls37* | *shl-1; shk-1; AWA::GFP* | Fig 4H, S3B, 5G |
| CX17101 | *slo-1(eg142); shk-1(ok1581); kyls37* | *slo-1; shk-1; AWA::GFP* | Fig 4H, S3B |
| CX17217 | *kcnl-2(ok2818); shk-1(ok1581); kyls37* | *kcnl-2; shk-1; AWA::GFP* | Fig S3B |
| CX17188 | *egl-2(rg4); shk-1(ok1581); kyls37* | *egl-2; shk-1; AWA::GFP* | Fig S3B |
| CX17290 | *kqt-3(tm0542); shk-1(ok1581); kyls37* | *kqt-3; shk-1; AWA::GFP* | Fig S3B |
| CX17258 | *slo-1(eg142); slo-2(nf101); shk-1(ok1581); kyls37* | *slo-1; slo-2; shk-1; AWA::GFP* | Fig 4H, S3B |
| CX17041 | *egl-19(ad695); kyls37* | *egl-19(gf); AWA::GFP* | Fig 6E-F |
| CX17462 | *kyEx6119[gpa-6::GCaMP6f 40ng + unc-122::dsRed 15ng]* | *AWA::GCaMP6f* | Fig 7A, B, F |
| CX17463 | *shk-1(ok1581); kyEx6120[gpa-6::GCaMP6f 40ng + unc-122::dsRed 15ng]* | *shk-1; AWA::GCaMP6f* | Fig 7C-E, G |

**Table S2.**
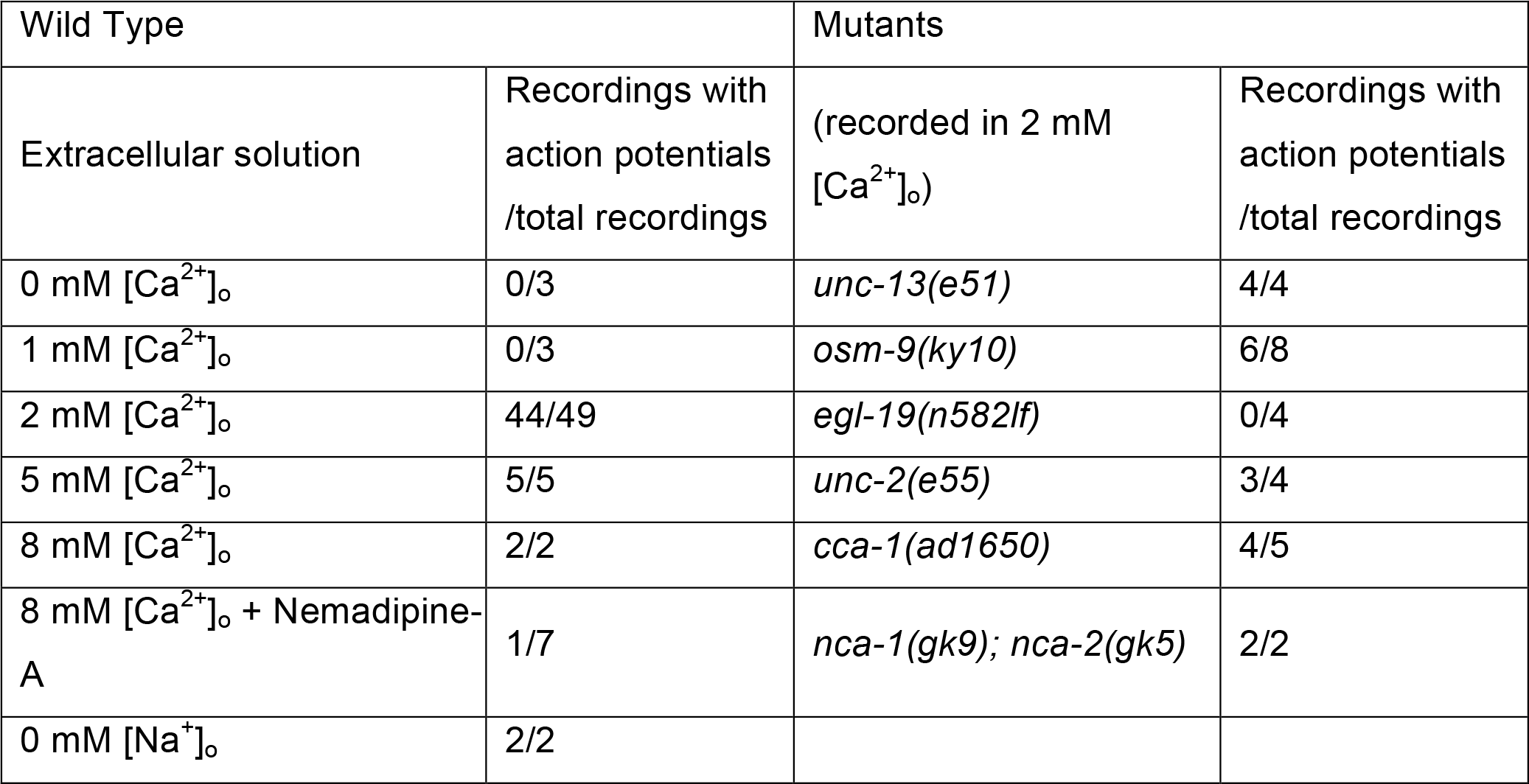
Effects of extracellular solutions and genetic mutations on spiking.

**Table S3.** AWA action potential features.

| Genotype or condition | n | Threshold |  | Amplitude |  | Width |  | AHP |  |
| --- | --- | --- | --- | --- | --- | --- | --- | --- | --- |
|  |  | Threshold (mV) | P vs. WT | Amplitude (mV) | P vs. WT | Width (ms) | P vs. WT | AHP (mV) | P vs. WT |
| WT | 149 | -34.65 ± 0.27 |  | 26.19 ± 0.40 |  | 35.93 ± 0.81 |  | 1.56 ± 0.26 |  |
| *slo-1* | 13 | -28.50 ± 0.92 | **** | 23.27 ± 1.33 | ns | 36.17 ± 2.43 | ns | 6.83 ± 1.18 | **** |
| *unc-2* | 13 | -30.39 ± 0.56 | **** | 23.75 ± 1.07 | ns | 45.89 ± 2.53 | ns | 3.58 ± 1.08 | ns |
| *cca-1* | 18 | -33.89 ± 0.44 | ns | 24.77 ± 1.25 | ns | 41.34 ± 2.44 | ns | 2.35 ± 0.69 | ns |
| WT + 4AP | 13 | -28.67 ± 0.96 | **** | 37.10 ± 1.65 | **** | 81.65 ± 3.58 | **** | 7.29 ± 0.92 | **** |
| *shk-1* | 41 | -33.30 ± 0.64 | ns | 36.17 ± 1.05 | **** | 139.24 ± 8.88 | **** | 6.82 ± 1.22 | **** |
| *shl-1* | 59 | -33.47 ± 0.33 | ns | 28.17 ± 0.76 | ns | 39.33 ± 1.55 | ns | 3.61 ± 0.43 | ** |
| *unc-13* | 12 | -32.36 ± 1.07 | ns | 22.18 ± 1.96 | ns | 35.95 ± 3.75 | ns | 4.75 ± 1.49 | ns |
| *slo-2* | 22 | -33.63 ± 0.40 | ns | 26.02 ± 1.13 | ns | 33.80 ± 1.22 | ns | 3.32 ± 0.78 | ns |
| *egl-19(gf)* | 24 | -36.52 ± 0.48 | ns | 24.24 ± 0.58 | ns | 34.02 ± 1.24 | ns | 5.77 ± 1.10 | **** |
| *nca-1;nca-2* | 24 | -31.73 ± 0.62 | *** | 23.32 ± 1.41 | ns | 38.33 ± 2.17 | ns | 3.13 ± 0.66 | ns |
| *osm-9* | 55 | -38.13 ± 0.35 | **** | 24.11 ± 0.73 | ns | 27.86 ± 1.26 | ns | 3.44 ± 0.44 | * |
| *shw-1* | 20 | -31.61 ± 0.71 | ** | 23.13 ± 0.67 | ns | 37.80 ± 1.97 | ns | 3.32 ± 0.61 | ns |
| *exp-3(gf)* | 14 | -31.78 ± 0.69 | * | 23.07 ± 1.60 | ns | 40.23 ± 2.07 | ns | 3.37 ± 0.72 | ns |
| *kcnl-2* | 32 | -29.47 ± 1.02 | **** | 23.79 ± 0.69 | ns | 35.79 ± 2.02 | ns | 3.75 ± 0.52 | * |
| WT 5mM [Ca<sup>2+</sup>]<sub>o</sub> | 20 | -30.05 ± 0.69 | **** | 34.23 ± 1.13 | **** | 18.98 ± 1.30 | ** | 7.04 ± 0.81 | **** |
| WT 8mM [Ca<sup>2+</sup>]<sub>o</sub> | 13 | -29.07 ± 0.65 | **** | 32.27 ± 1.27 | *** | 20.86 ± 1.03 | ns | 4.62 ± 0.32 | ns |
| *exp-2* | 4 | -29.56 ± 1.37 |  | 19.09 ± 1.65 |  | 35.65 ± 0.38 |  | 6.21 ± 1.93 |  |
| *egl-2* | 6 | -29.74 ± 1.04 |  | 27.25 ± 1.13 |  | 27.10 ± 1.34 |  | 4.67 ± 1.25 |  |
| *shw-2* | 5 | -31.63 ± 0.80 |  | 19.96 ± 1.31 |  | 48.00 ± 3.02 |  | 3.69 ± 1.11 |  |
|  |  | Threshold (mV) | P vs. *shk-1* | Amplitude (mV) | P vs. *shk-1* | Width (ms) | P vs. *shk-1* | AHP (mV) | P vs. *shk-1* |
| *shk-1* | 41 | -33.30 ± 0.64 |  | 36.17 ± 1.05 |  | 139.24 ± 8.88] |  | 6.82 ± 1.22 |  |
| *slo-1;shk-1* | 18 | -31.35 ± 0.86 | ns | 40.25 v 1.31 | ns | 141.04 ± 10.09] | ns | 10.90 [1.12 | ns |
| *slo-2;shk-1* | 39 | -30.15 ± 0.75 | ** | 35.82 ± 1.16 | ns | 135.24 ± 13.34] | ns | 8.87 ± 2.18 | ns |
| *kcnl-2;shk-1* | 24 | -29.62 ± 0.64 | ** | 37.65 ± 1.00 | ns | 159.29 ± 15.90] | ns | 8.27 ± 2.18 | ns |
| *slo-1;slo-2;shk-1* | 8 | -26.98 ± 1.15 | *** | 36.47 ± 2.02 | ns | 132.88 ± 5.86] | ns | 13.02 ± 1.38 | ns |
| *shl-1;shk-1* | 18 | -28.93 ± 0.63 | ** | 39.42 ± 1.18 | ns | 137.24 v 13.49] | ns | 7.47 ± 1.18 | ns |
| *kqt-3;shk-1* | 28 | -31.48 ± 0.88 | ns | 36.44 ± 1.14 | ns | 117.11 v 6.79] | ns | 10.64 ± 0.94 | ns |
| *egl-2;shk-1* | 12 | -33.99 ± 1.27 | ns | 38.89 ± 1.77 | ns | 119.08 ± 16.14] | ns | 11.45 ± 1.97 | ns |
| WT + 4AP | 13 | -28.67 ± 0.96 | ** | 37.10 ± 1.65 | ns | 81.65 v 3.58] | * | 7.29 ± 0.92 | ns |

Results are presented as mean with standard error of the mean. n, number of action potentials. Data from all individual spikes were analyzed using Dunnett's test for multiple comparisons, and were not calculated for three genotypes with small n. \*\*\*\*, P<0.0001; \*\*\*, P<0.001; \*\*P<0.01; \*P<0.05; ns, not significant. Many mutations slightly affected the threshold and AHP; *shk-1* uniquely affected spike amplitude and width, resembling pharmacological administration of 4AP. Single spikes (Fig 2B,C) have a narrower width and larger AHP than spikes in burst.

**Table S4.** AWA Model Parameters.

| Symbol | Description | Value (WT/shk-1) |
| --- | --- | --- |
| $C$ | Membrane capacitance | 1.5 pF |
| $g_{Ca}$ | Calcium conductance | 0.1/0.13 nS |
| $v_{Ca}$ | Calcium reversal potential | 120 mV |
| $f$ | Calcium inactivation factor | 0.4 |
| $\tau_{c1}$ | Calcium channel activation time scale | 1.0 ms |
| $\tau_{c2}$ | Calcium channel inactivation time scale | 160.0 ms |
| $v_{c1}$ | Calcium channel activation voltage | -21.6 mV |
| $\Delta v_{c1}$ | Calcium channel activation slope | 9.17 mV |
| $v_{c2}$ | Calcium channel inactivation voltage | 16.2 mV |
| $\Delta v_{c2}$ | Calcium channel inactivation slope | -16.1 mV |
| $v_{c3}$ | Calcium channel inactivation time voltage scale | 23.0 mV |
| $\Delta v_{c3}$ | Calcium channel inactivation time voltage slope | 24.0 mV |
| $c_0$ | Calcium channel steady state normalization | 0.55 |
| $v_K$ | Potassium reversal potential | -84 mV |
| $g_{K1}$ | SHK-1 conductance | 1.5/0.0 nS |
| $v_{K1}$ | SHK-1 activation voltage | 2.0 mV |
| $\Delta v_{K1}$ | SHK-1 activation slope | 10.0 mV |
| $\tau_{K1}$ | SHK-1 switching time | 30.0 ms |
| $g_{K2}$ | First high-v potassium conductance | 1.0/2.0 nS |
| $v_{K2}$ | First high-v potassium channel activation voltage | 13.0 mV |
| $\Delta v_{K2}$ | First high-v potassium channel activation slope | 20.0 mV |
| $\tau_{K2}$ | First high-v potassium channel switching time | 1000 ms |
| $g_{K3}$ | Second high-v potassium conductance | 0.1 nS |
| $v_{K3}$ | Second high-v potassium channel activation voltage | -25.0 mV |
| $\Delta v_{K3}$ | Second high-v potassium channel activation slope | 5.0 mV |
| $\tau_{K3}$ | Second high-v potassium channel switching time | 1000 ms |
| $g_{Kr}$ | Slowly inactivating (Kr) K conductance | 0.39/0.77 nS |
| $v_{Kr1}$ | Slowly inactivating (Kr) K-channel activation voltage | -42.0 mV |
| $\Delta v_{Kr1}$ | Slowly inactivating (Kr) K-channel activation slope | 5.0 mV |
| $v_{Kr2}$ | Slowly inactivating (Kr) K-channel inactivation voltage | -42.0 mV |
| $\Delta v_{Kr2}$ | Slowly inactivating (Kr) K-channel inactivation slope | 5.0 mV |
| $\tau_{Kr}$ | Slowly inactivating (Kr) K-channel inactivation time | 1800 ms |
| $g_{Ka}$ | Slowly activating (Kr<sub>a</sub>) K-channel conductance | 0.91/1.98 nS |
| $v_{Ka1}$ | Slowly activating (Kr<sub>a</sub>) K-channel activation voltage | -52.0 mV |
| $\Delta v_{Ka1}$ | Slowly activating (Kr<sub>a</sub>) K-channel activation slope | 20.0 mV |
| $v_{Ka2}$ | Slowly activating (Kr<sub>a</sub>) K-channel time voltage scale | -32.0 mV |
| $\Delta v_{Ka2}$ | Slowly activating (Kr<sub>a</sub>) K-channel time voltage slope | 4.0 mV |
| $\tau_{Ka1}$ | Slowly activating (Kr<sub>a</sub>) K-channel fast activation time | 27000 ms |
| $\tau_{Ka2}$ | Slowly activating (Kr<sub>a</sub>) K-channel slow activation time | 3000 ms |
| $g_{Kir}$ | Inward rectifying potassium channel conductance | 0.8 nS |
| $I_0$ | Inward rectifying potassium channel saturated current | 5.0 pA |
| $\beta$ | Inward rectifying potassium channel smoothing factor | 5.0 |
| $g_L$ | Leak current conductance | 0.25 nS |
| $v_L$ | Leak current reversal potential | -65.0 mV |

**Supplemental Figure 1.**
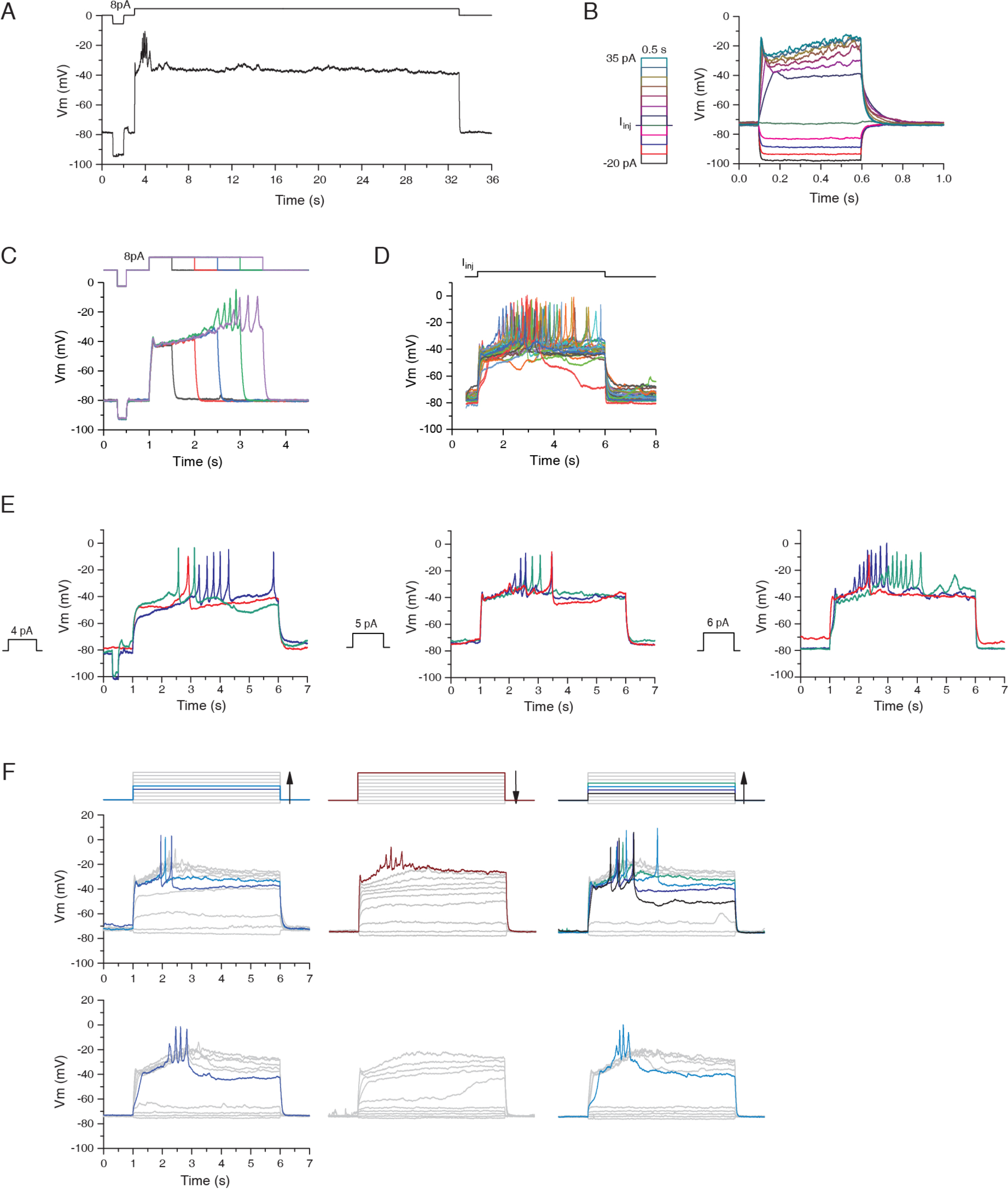
Additional properties of action potentials in AWA. **A.** Representative WT AWA current-clamp trace with a single 30 s-long current injection at 8 pA. A short burst of action potentials was induced in the first 2 seconds after stimulus onset. Membrane potential remained static at the plateau state for the rest of the stimulation. **B.** Representative WT AWA current-clamp traces under a series of 0.5 s current injections of variable amplitudes. Action potentials were not induced by short stimuli for any amplitude of current injected. **C.** Representative WT AWA current-clamp traces under a series of current injections at 8 pA of varying durations. 0.5 s (black), 1 s (red) and 1.5 s (blue) current injection steps failed to induce action potentials in this example. Longer stimuli at either 2 s (green) or 2.5 s (purple) induced action potential bursts. **D.** Overlay of action potential traces from 27 WT AWA preps showing the stochastic nature of the spikes. The spike time distribution of all the action potentials for this set of data is shown in Fig 2E. **E.** Representative action potential traces induced by current injection at 4, 5 or 6 pA showing variable timing and number of spikes induced by a given stimulus. **F.** Representative WT AWA traces under a series of current injections at 2 pA per step with opposite execution sequence. When the current injection series started from hyperpolarization steps, action potentials were induced in both cases (first column). Immediately following the first current injection series, the series was executed in the opposite direction and no action potentials were induced in the same cell (middle column), except for the partial spikes in the top trace (black trace). The third column shows recovery of action potential firing when the current injection series was repeated from a hyperpolarizing direction.

**Supplemental Figure 2.**
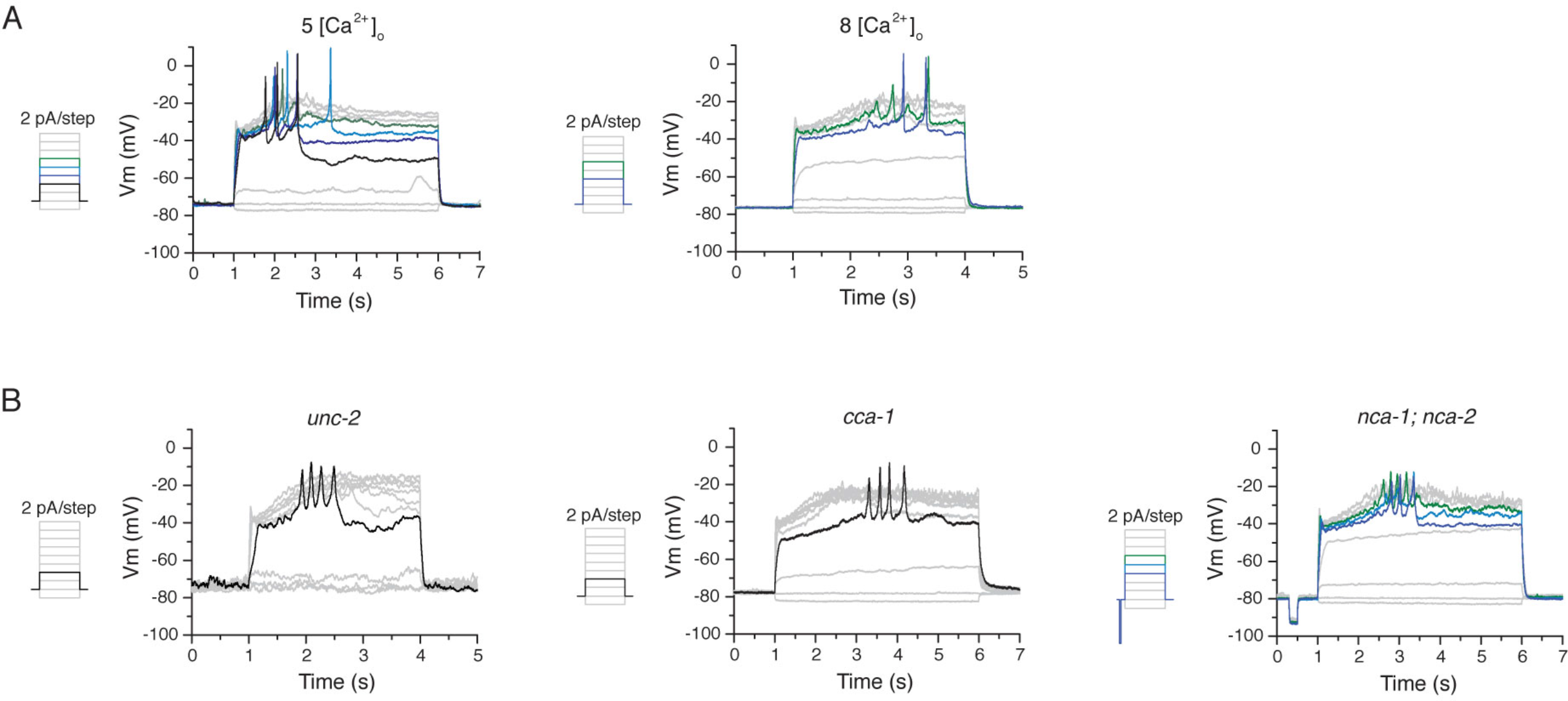
Spiking in different extracellular calcium concentrations and calcium channel mutants. **A.** Representative current-clamp traces recorded from WT AWA in extracellular solutions with 5 or 8 mM [Ca^2+^]_o_. **B.** Representative current-clamp traces recorded from calcium channel mutants: CaV2 *(unc-2)*, CaV3 *(cca-1)*, or NALCN *(nca-1;nca-2)*.

**Supplemental Figure 3.**
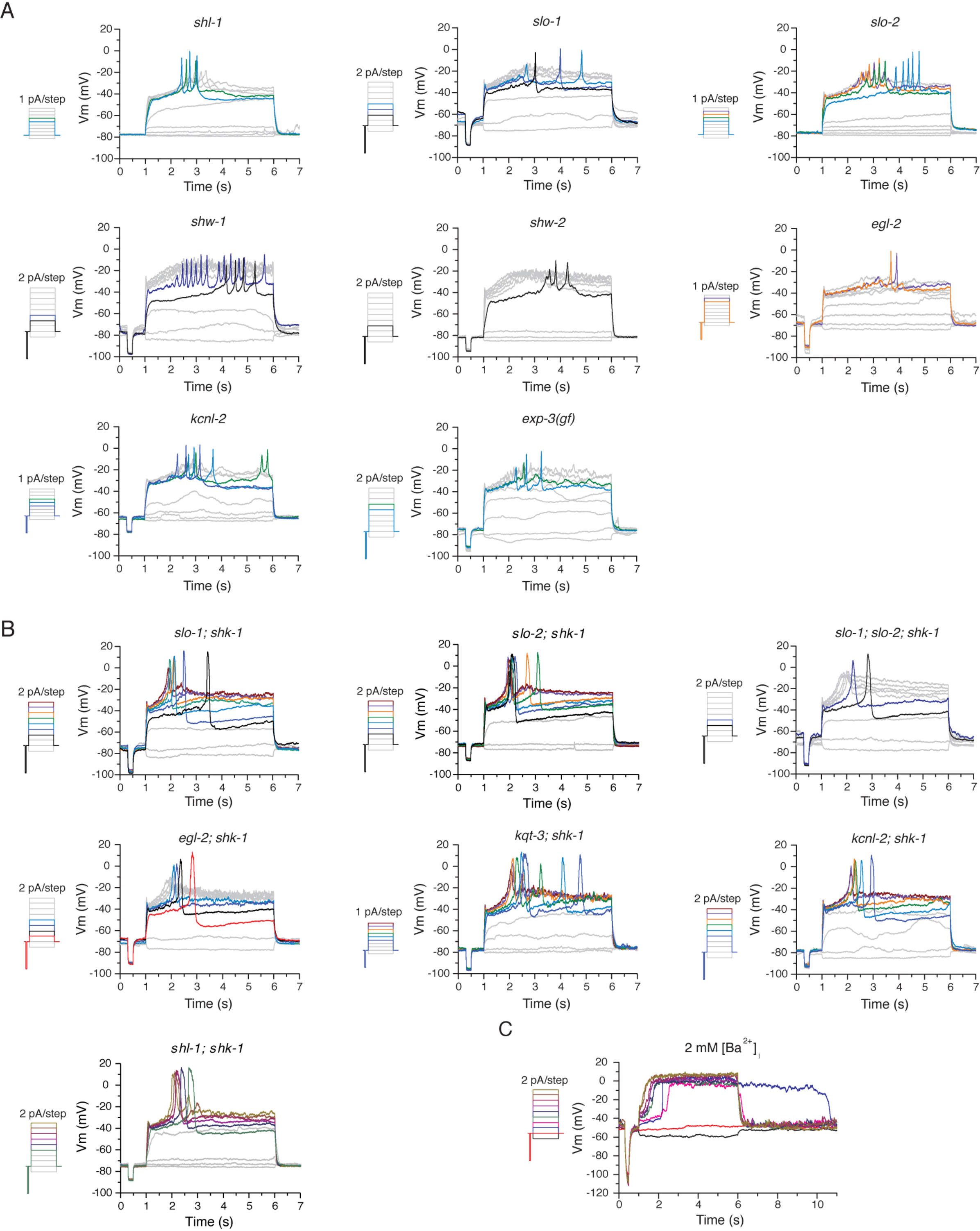
Spiking in potassium channel mutants. **A.** Representative AWA traces recorded from various K channel mutants under a series of current injection steps with 1 pA or 2 pA increments. Traces with spiking activities are in color. Non-spiking traces are in gray. **B.** Representative AWA traces recorded from double mutants of *shk-1* and other K channels under a series of current injection steps with 1 pA or 2 pA increments. **C.** Representative AWA membrane potential recording with 2 mM extracellular Ca^2+^ substituted with Ba^2+^ under a series of current injection steps at 2 pA increments. The fast depolarization under each positive current injection step is equivalent of the upstroke of action potentials recorded in Ca^2+^ containing solutions. However, repolarization did not occur until the end of the stimuli, and sometimes the depolarization lasted longer beyond the stimulation pulse, as in the blue trace in this example.

**Supplemental Figure 4.**
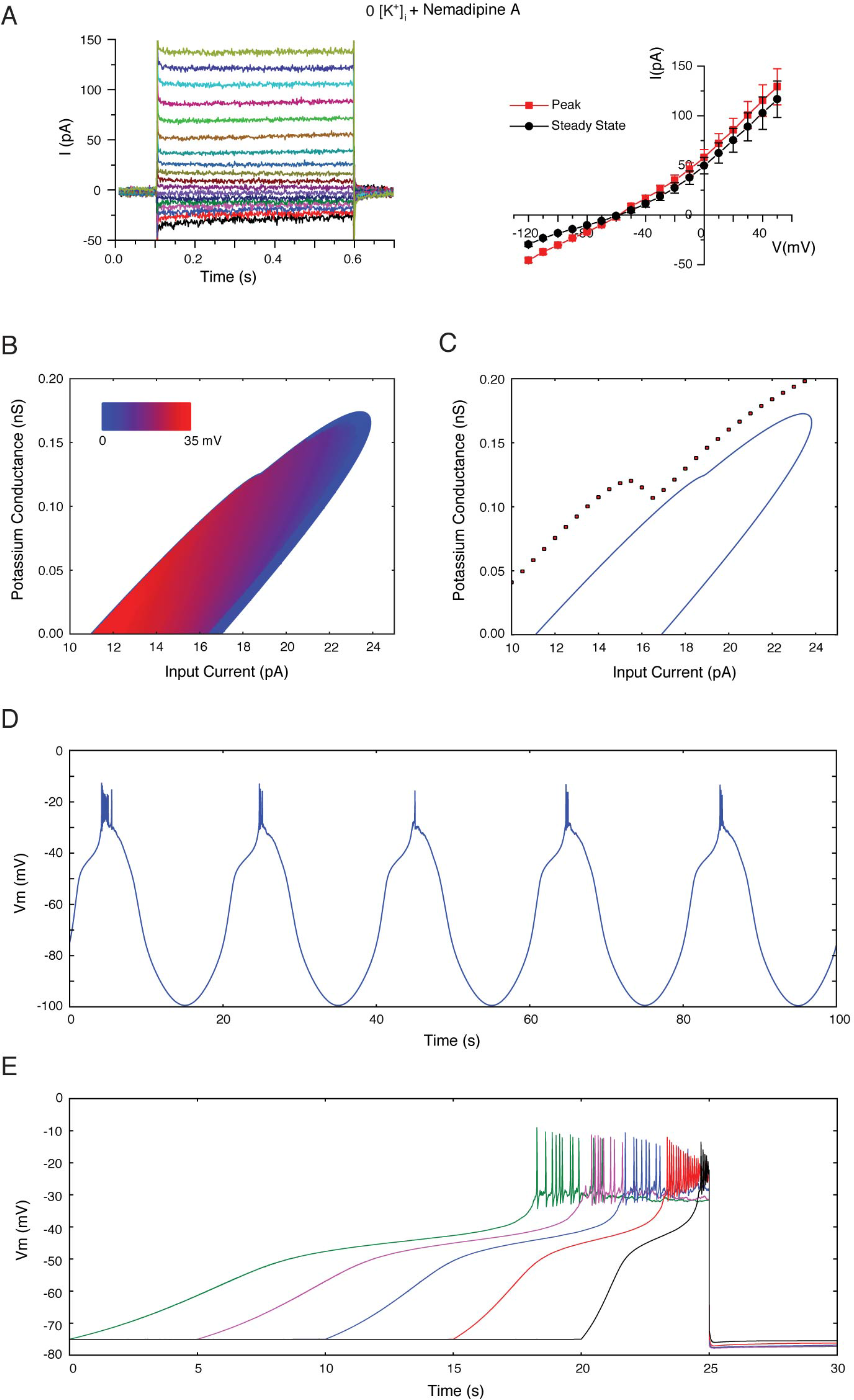
Design of the mathematical model. **A.** Left: Representative whole-cell currents in WT AWA recorded under voltage-clamp in conditions as in Fig 5A, but with 10 uM Nemadipine-A added to the extracellular solution. Note the absence of the small peak depolarizing currents seen in Fig 5A. Right: I-V relationships for peak and steady-state currents under this condition (n = 4). **B.** Bifurcation diagram of the model in terms of input current and added potassium conductance. The colored region is bounded by a line of Hopf bifurcations and marks the values of *I* and *g*_K_ where spiking will occur. The color indicates the amplitude of spiking at each point in the *I*-*g*_K_ plane. **C.** The same bifurcation diagram as in B, with black dots showing the minimum value of the potassium conductance from the combined Kr and Kr_a_ channels during a 5 second current pulse of the indicated magnitude. **D.** Simulation of the model with a sinusoidally varying current input, at 0.05 Hz and 22 pA amplitude. **E.** Simulation of the model with a linearly increasing current input according to the same protocol used in Fig 8B, except with final current equal to 14 pA.

**Supplemental Figure 5.**
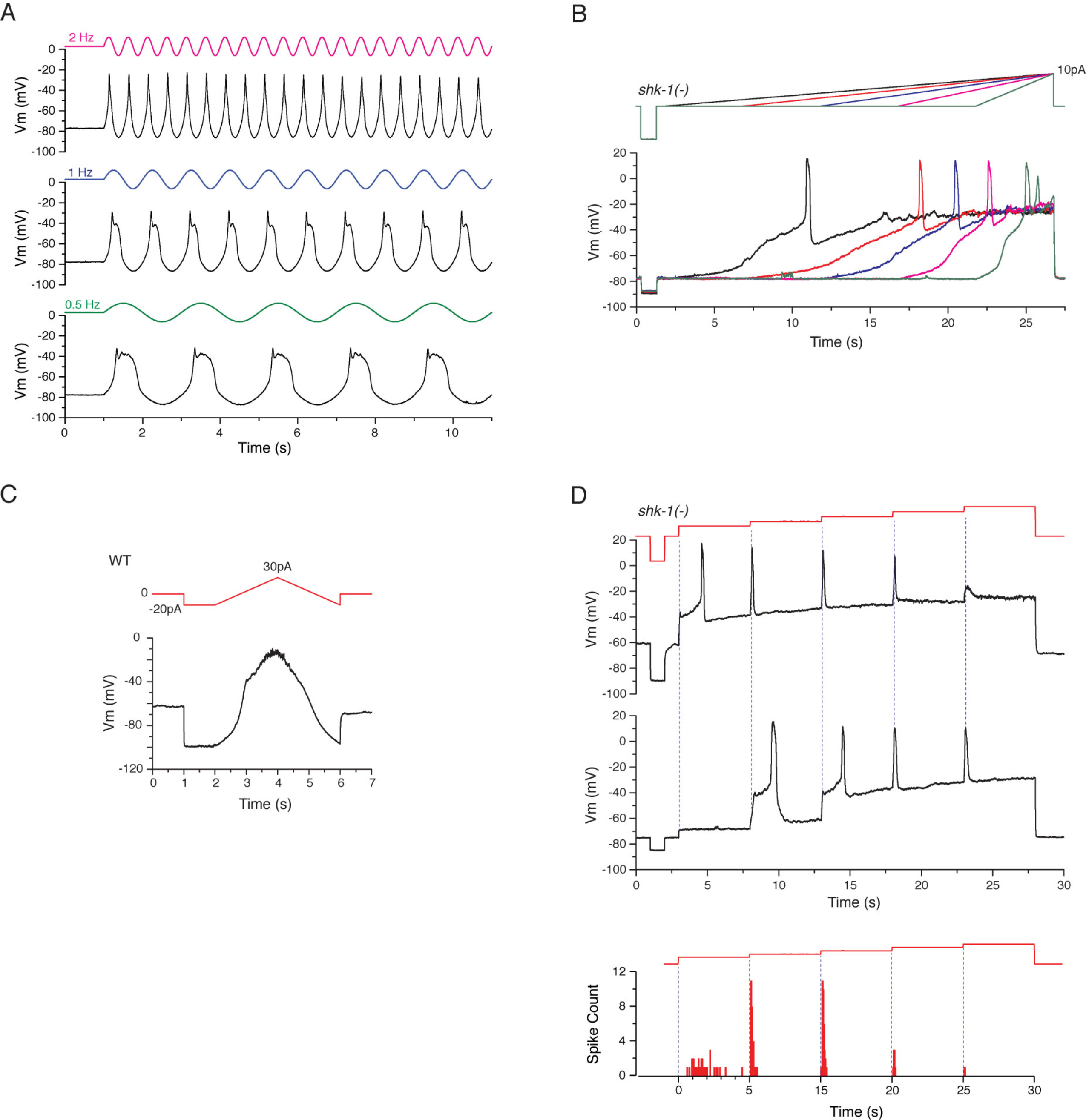
AWA action potentials may encode specific stimulus features. **A.** Representative current-clamp traces recorded from WT AWA showing action potentials were not induced by high frequency sinusoidal stimuli (2 Hz in magenta, 1 Hz in blue and 0.5 Hz in green). **B.** Representative AWA action potentials evoked by current injection ramps at different slopes in *shk-1; kcnl-2* mutant (black: 10 pA/25 s; red: 10 pA/20 s; blue: 10 pA/15 s; pink: 10 pA/10 s; green: 10 pA/5 s). **C.** AWA membrane potential recording trace under a current injection ramp protocol at 50 pA/2 s. There was a fast transition in membrane potential between the resting state and the plateau state, but action potentials were not induced by the fast ramp stimulus. **D.** Two examples of AWA recordings with action potentials induced by a five-step current injection. Each step is 5 s in duration. The upper trace is an AWA recording from a *shk-1; shl-1* mutant. The first 4 pA step induced a single spike with a ~2 s delay and several subsequent 2 pA steps induced single instantaneous spikes. The middle trace is an AWA recording from a *shk-1; slo-2* mutants. The first 4 pA step did not induce action potentials or depolarize AWA’s membrane potential sufficiently to reach the plateau state. The second 2 pA step induced a single spike with a ~2 s delay, while the AHP brought the membrane potential back to the resting state. The third 2 pA step induced another spike with a delay and maintained the membrane potential at the plateau state. The last two steps induced spikes without delays. Lower panel: spike time distribution of all spikes recorded in AWA from strains with *shk-1* mutations under the 5-step current injection protocol. Several single and double mutants strains with *shk-1* mutations were pooled together for this analysis; action potentials in AWA behave similarly in those strains (Fig 4H and Fig S3B). Bin size is 50 ms. 1st step: n = 32; 2nd step: n = 39; 3rd step: n = 35; 4th step: n = 8; 5th step: n = 2. Blue dotted lines indicate the onset of each upstep. The unusual firing pattern shown in the middle panel is not included. In all other cases, the first step induced action potential spikes with variable delays, and subsequent upsteps induced spikes from the plateau state without delays.

**Supplemental Figure 6.**
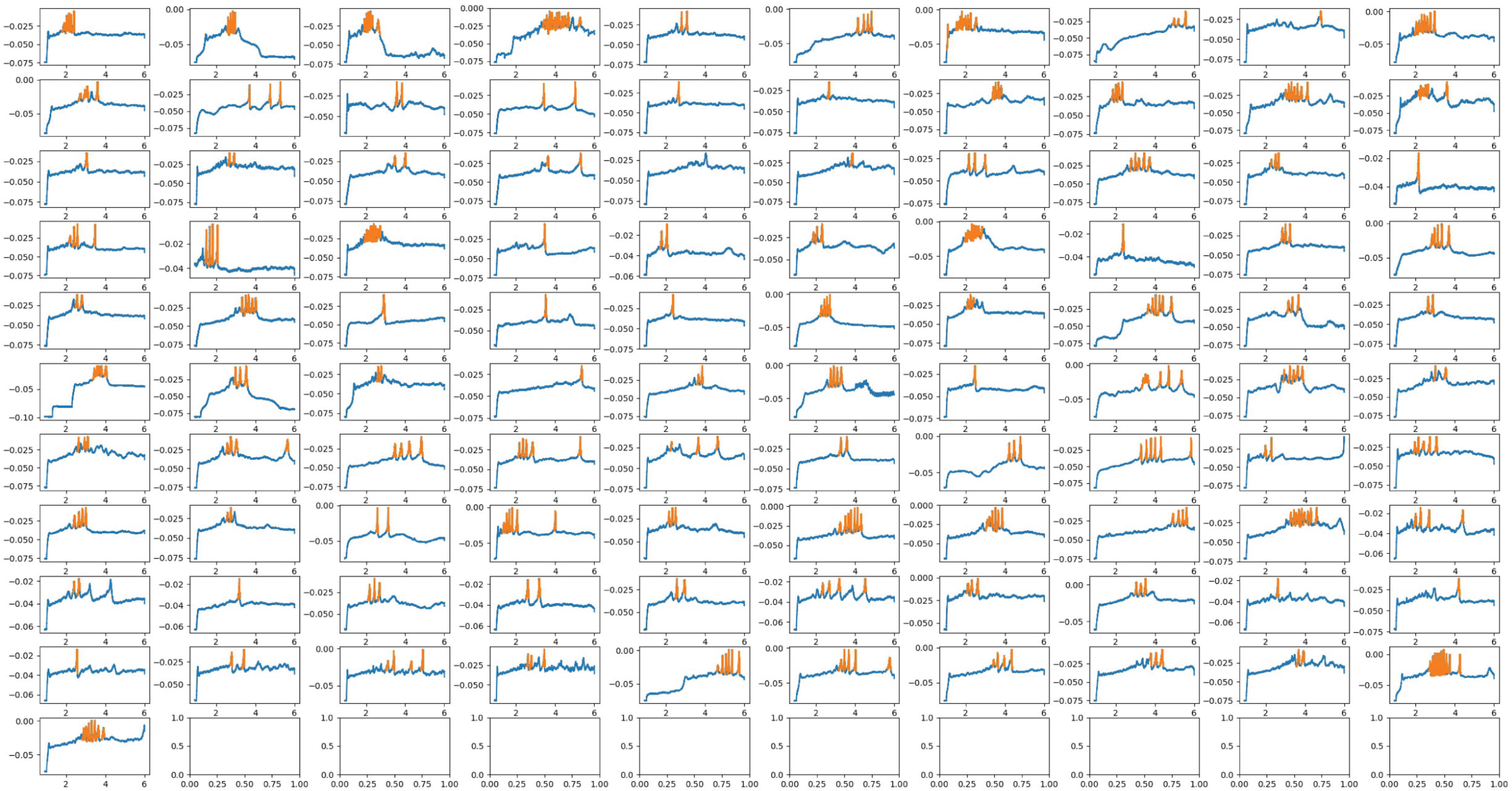
All algorithm-detected WT action potential spikes used for analysis. All action potential spikes identified by our spike-detection algorithm from WT AWA recordings. Y axis: voltage (V). X axis: time (s).

**Supplemental Figure 7.**
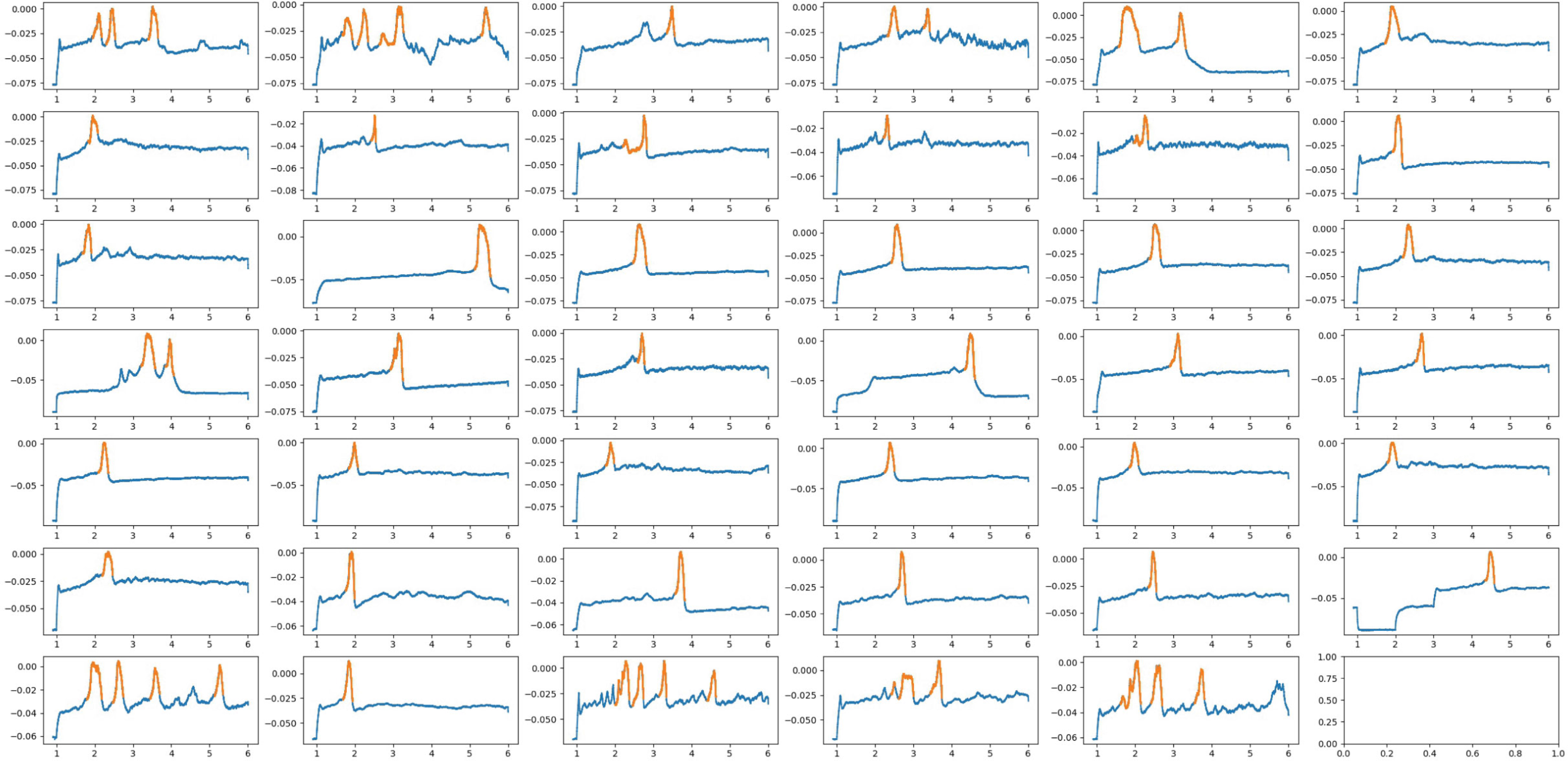
All algorithm-detected shk-1 action potential spikes used for analysis. All action potential spikes identified by our spike-detection algorithm from *shk-1* AWA recordings. Y axis: voltage (V). X axis: time (s).

